# NADPH consumption by L-cystine reduction creates a metabolic vulnerability upon glucose deprivation

**DOI:** 10.1101/733162

**Authors:** James H. Joly, Alireza Delfarah, Philip S. Phung, Sydney Parrish, Nicholas A. Graham

## Abstract

The consequences of metabolic reprogramming in cancer can include an increased dependence on metabolic substrates such as glucose for survival. As such, the vulnerability of cancer cells to glucose deprivation creates an attractive opportunity for therapeutic intervention. Because it is not possible to starve tumors of glucose *in vivo*, we sought to identify the mechanisms regulating cancer cell death upon glucose deprivation and then design combinations of inhibitors to mimic glucose deprivation-induced cell death. Using metabolomic profiling, we found that cells undergoing glucose deprivation-induced cell death exhibited dramatic accumulation of intracellular L-cysteine and its oxidized dimer, L-cystine, and depletion of the antioxidant glutathione. Building on this observation, we show that glucose deprivation-induced cell death is driven not by lack of glucose but rather by L-cystine import. Following glucose deprivation, the import of L-cystine and subsequent reduction to L-cysteine depleted both NADPH and glutathione, thereby allowing toxic accumulation of reactive oxygen species. Consistent with this model, we found that the glutamate/cystine antiporter, xCT, was required for sensitivity to glucose deprivation. We searched for glycolytic enzymes whose expression is essential for survival of cancer cells with high xCT expression and identified the glucose transporter GLUT1. We therefore tested a drug combination co-targeting GLUT1 and glutathione synthesis and found that these drugs induced synthetic lethal cell death in high xCT-expressing cell lines susceptible to glucose deprivation. These results indicate that co-targeting GLUT1 and glutathione synthesis is a potential therapeutic approach in tumors dependent on glucose for survival.

## Introduction

Cancer cells exhibit an altered metabolism compared to non-transformed cells, consuming glucose and producing lactate at much higher rates than non-transformed cells. The re-wiring of metabolic networks in cancer cells is thought to help satisfy the extreme biosynthetic demands required of highly proliferative cells, including synthesis of lipids, proteins, and nucleic acids. This differential metabolism suggests that targeting metabolic dependencies specific to cancer cells could be an effective approach for cancer therapy^1–5^.

One consequence of metabolic reprogramming is that some cancer cells become dependent on metabolic substrates including glucose and glutamine for survival^6, 7^. This glucose or glutamine “addiction” has been linked to signaling through AKT and AMPK^8–,10^, mTORC1^11^, and tyrosine kinases^12^. Other studies have linked glucose deprivation-induced death to mechanisms independent of glycolysis including Ca^2+^ influxes through the L-type calcium channel Ca_v_1.3^13^ and the glutamate/cystine antiporter xCT (*SLC7A11*)^14–16^. However, therapeutically targeting the glucose addiction phenotype remains a challenge^2^ because 1) tumors cannot be completely starved of glucose *in vivo*; and 2) cancer cells exhibit a spectrum of sensitivity to glucose deprivation^12^. In fact, individual inhibitors of glucose metabolism alone have achieved limited success^17–20^, perhaps because other organs also exhibit high glucose uptake including brain and heart^21^.

A promising trend in cancer therapy is the concept of synthetic lethality, whereby cell death occurs following the simultaneous inactivation of multiple gene products but not the inactivation of either gene product alone^22^. We reasoned that combinations of inhibitors mimicking glucose deprivation-induced cell death might exhibit synthetic lethal interactions specifically in cancer cells susceptible to glucose deprivation. We thus sought to identify the mechanisms regulating cancer cell death following glucose deprivation. Using metabolomic profiling of sensitive and insensitive cancer cell lines, we found that glucose deprivation kills vulnerable cancer cells due to a redox imbalance driven by L-cystine import and subsequent consumption of NADPH during reduction to L-cysteine. In addition, we identified that expression of the specific light chain of the glutamate/cystine antiporter, xCT, is predictive of sensitivity to glucose deprivation. Building upon these findings, combined inhibition of glucose transporter 1 (GLUT1) and the rate limiting step of glutathione synthesis, glutamate-cysteine ligase (GCL), induced a synthetic lethal interaction in glucose deprivation-sensitive cancer cells. Finally, we identified a subset of glioblastoma tumors that exhibit high xCT expression compared to non-malignant cells. Taken together, our results define a new synthetic lethal combination therapy that exploits the glucose addiction phenotype by mimicking glucose deprivation-induced cell death even in the presence of glucose.

## Results

### Metabolomics identifies accumulation of L-cystine and L-cysteine and depletion of glutathione as markers of glucose deprivation sensitivity

To investigate the metabolic profile of glucose deprivation-induced cell death, we first tested the response of a panel of six GBM and two sarcoma cell lines to glucose deprivation. Among the GBM cell lines, we found that A172 exhibited strong resistance, LN229 and U118MG exhibited medium sensitivity, and LN18, T98, and U87 exhibited high sensitivity to glucose deprivation (Fig. 1A)^23, 24^. For the sarcoma cell lines, HT161 and TC32 exhibited resistance and sensitivity, respectfully, to glucose deprivation (Supporting Fig. 1A). We then used liquid chromatography-mass spectrometry (LC-MS) metabolomics to measure changes in metabolite abundance and TCA cycle flux upon glucose deprivation^25–27^. Using two sensitive cell lines (LN18, T98) and one medium resistant cell line (LN229), we quantified levels of 97 polar metabolites in cells cultured with or without glucose for 3 h (Supporting Table 1). To identify metabolites correlated with glucose deprivation sensitivity, we ranked metabolite abundances by a ratio-of-ratios comparing sensitive to resistant cells (Fig. 1B). This metric identified L-cystine (the oxidized dimer of L-cysteine, rank #1), L-cysteine (#2), and reduced glutathione (GSH) (#97) as the most differentially-changed metabolites between sensitive and resistant cells. In addition, the GSH precursor γ-glutamylcysteine (γ-GCS) was also differentially regulated between sensitive and resistant cells (rank 93 of 97).

**Figure 1.**
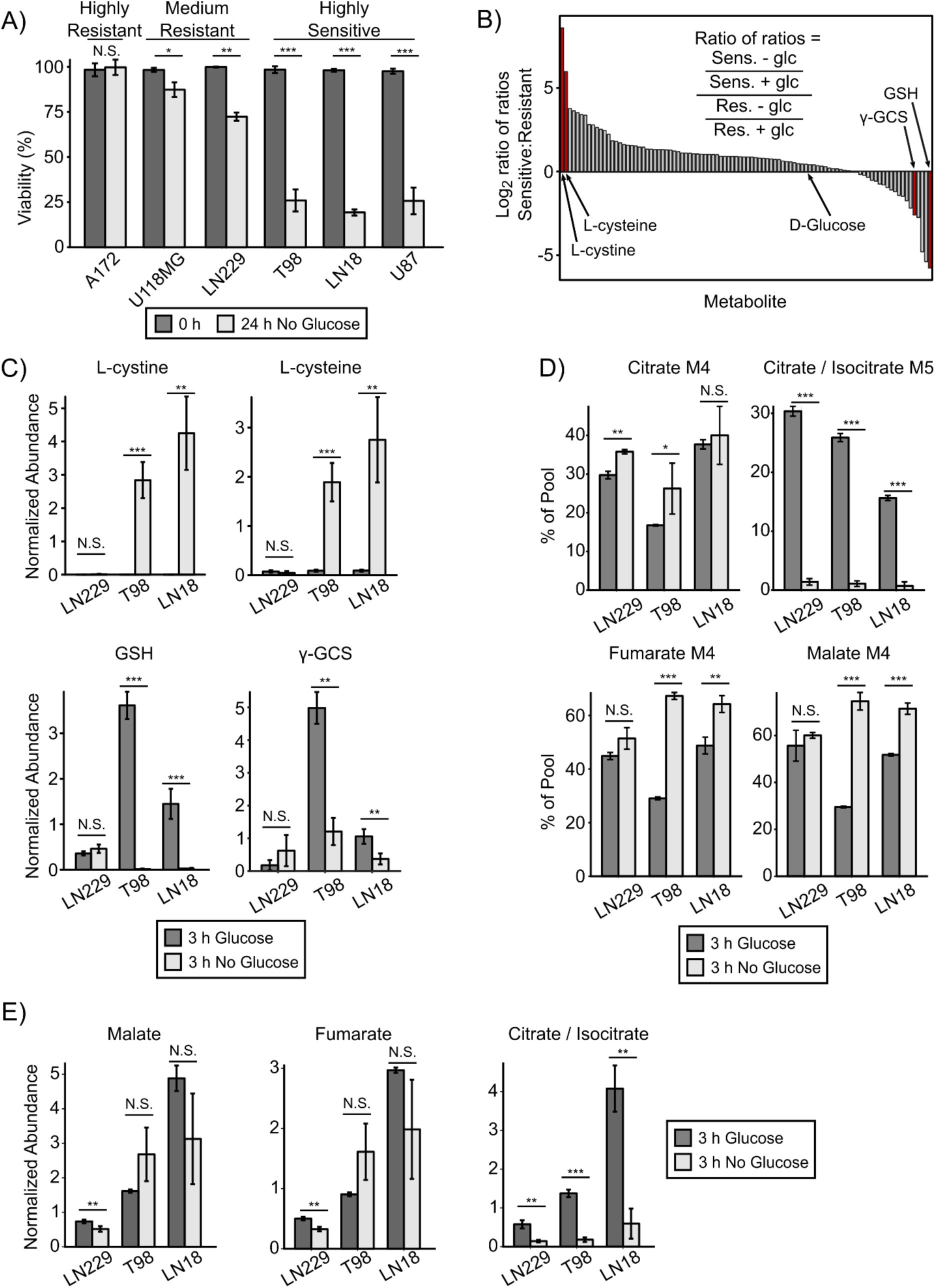
Accumulation of L-cystine and L-cysteine and depletion of reduced glutathione are metabolic markers of sensitivity to glucose deprivation. **A)** A panel of GBM cell lines was subjected to glucose deprivation for 24 h, and viability was measured by trypan blue exclusion. A172 and U118MG exhibited strong resistance, LN229 cells exhibited medium resistance, and T98, LN18, and U87 cells exhibited high sensitivity to glucose deprivation. N.S. denotes p > 0.05, * denotes p < 0.05, ** denotes p < 0.01, *** denotes p < 0.001 by Student’s t-test (n=3). **B)** Two glucose deprivation-sensitive cell lines (T98, LN18) and one medium resistant cell line (LN229) were deprived of glucose for 3 h in the presence of L-[U-^13^C]-Glutamine and then profiled by LC-MS metabolomics. A total of 97 metabolites were identified and quantified. Metabolite pool sizes were ranked using a log_2_ ratio-of-ratios metric ([Average of T98 −/+ glucose and LN18 −/+ glucose] / [LN229 −/+ glucose]). L-cystine and L-cysteine, and reduced glutathione (GSH) were the most differentially regulated metabolites (red bars). γ-glutamylcysteine (γ-GCS) was also differentially regulated in glucose deprivation-sensitive cells (rank 93 of 97). **C)** Bar plots from LC-MS metabolomics in B) showing that L-cystine and L-cysteine were accumulated, and GSH and γ-GCS were depleted in glucose deprivation-sensitive GBM cells following 3 h of glucose deprivation. * denotes p < 0.05, ** denotes p < 0.01, *** denotes p < 0.001 by Student’s t-test (n=3). **D)** Metabolite isotopomer distributions for selected TCA cycle metabolites after 3 h with or without glucose in the presence of L-[U-^13^C]-Glutamine. Glucose deprivation inhibited reductive carboxylation of α-ketoglutarate as indicated by decreased percentage of citrate M5. Forward flux through the TCA cycle was increased in glucose deprivation-sensitive cells, as indicated by increased percentages of fumarate M4 and malate M4. N.S. denotes p > 0.05, * denotes p < 0.05, ** denotes p < 0.01, *** denotes p < 0.001 by Student’s t-test (n=3). **E)** Bar plots from LC-MS metabolomics in B) showing the response of malate, fumarate, and citrate pool sizes to 3 h of glucose deprivation. Malate and fumarate levels were not significantly affected, whereas citrate levels were substantially decreased in all three cell lines. There was no consistent trend for any of the three metabolites with sensitivity to glucose deprivation. N.S. denotes p > 0.05, * denotes p < 0.05, ** denotes p < 0.01, *** denotes p < 0.001 by Student’s t-test (n=3).

Examination of these differentially-regulated metabolites revealed that the glucose deprivation-sensitive T98 and LN18 cells exhibited 800- and 30-fold accumulation of L-cystine and L-cysteine, respectively (Fig. 1C). In contrast, reduced GSH was completely depleted and γ-GCS levels dropped by ∼80% following glucose deprivation in sensitive T98 and LN18 cells (Fig. 1C). In contrast, LN229 cells exhibited little to no change in levels of these metabolites. Next, we used a targeted LC-MS metabolomics assay to confirm these trends in one additional sensitive cell line (U87) and one additional resistant cell line (A172). Indeed, we corroborated that sensitive U87 cells accumulated L-cystine and L-cysteine and depleted GSH following 3 h of glucose deprivation, whereas the highly resistant cell line A172 showed no significant changes in these metabolites (Supporting Fig. 1B). Together, these data demonstrate that glucose deprivation sensitive cells exhibit a metabolic signature of L-cystine and L-cysteine accumulation and GSH depletion upon glucose deprivation.

We next investigated changes in TCA cycle flux upon glucose deprivation by incubating cells with L-[U-^13^C]-glutamine in the presence or absence of glucose. We found that >90% of glutamine was labeled in 3 h in both resistant (LN229) and sensitive (LN18, T98) cell lines (Supporting Fig. 1C and Supporting Table 2 and Supporting Table 3). Examination of the isotopomer distributions for TCA cycle metabolites showed a significant decrease in the percentage of citrate M5 upon glucose deprivation, indicating decreased reductive carboxylation flux from α-ketoglutarate to citrate in glucose-deprived cells^28, 29^ (Fig. 1D). In contrast, forward flux through the TCA cycle was increased upon glucose deprivation in sensitive but not resistant cell lines, as indicated by increased labeling of malate M4 and fumarate M4 upon glucose deprivation (Fig. 1D). Interestingly, both sensitive and resistant cell lines exhibited increased labeling of L-aspartate M4 upon glucose starvation, further suggesting increased forward flux through the TCA cycle (Supporting Fig. 1C). Malate and fumarate pool sizes were not consistently affected by glucose deprivation in either direction, whereas citrate and aspartate levels were substantially decreased and increased, respectively, in all three cell lines (Fig. 1E and Supporting Fig. 1D). Taken together, these data demonstrate that cancer cells shift towards oxidative, rather than reductive, glutamine metabolism when deprived of glucose.

### Glutamate-cysteine ligase (GCL) activity moderately regulates resistance to glucose deprivation

The differential responses of L-cystine, L-cysteine, γ-GCS, and GSH upon glucose deprivation is particularly interesting because these four molecules all belong to the GSH synthesis pathway. GSH is a tripeptide formed by sequential ligation of the amino acids L-cysteine, L-glutamate, and glycine, and is the most abundant antioxidant in the cell^30, 31^. L-glutamate and L-cysteine are first ligated to form γ-GCS, and then γ-GCS and glycine are joined to form GSH. We therefore examined the levels of other GSH synthesis precursors in response to glucose deprivation. In contrast to L-cysteine, the levels of L-glutamate and glycine were unchanged upon glucose deprivation in both sensitive and resistant cells (Fig. 2A). Additionally, levels of oxidized glutathione (GSSG) and ATP, which is required for both γ-GCS and GSH formation, remained abundant following glucose deprivation in all cell lines. Overlaying the quantitative changes in metabolite pool size on a pathway diagram of GSH synthesis suggested that glucose deprivation-sensitive cells are unable to synthesize GSH despite the presence of abundant synthetic precursors (Fig. 2B and Supporting Fig. 2A). Together, this data suggested that glucose deprivation-sensitive cells may exhibit inhibition of GSH synthesis following glucose deprivation at the enzymatic step that ligates L-glutamate and L-cysteine to form γ-GCS.

**Figure 2.**
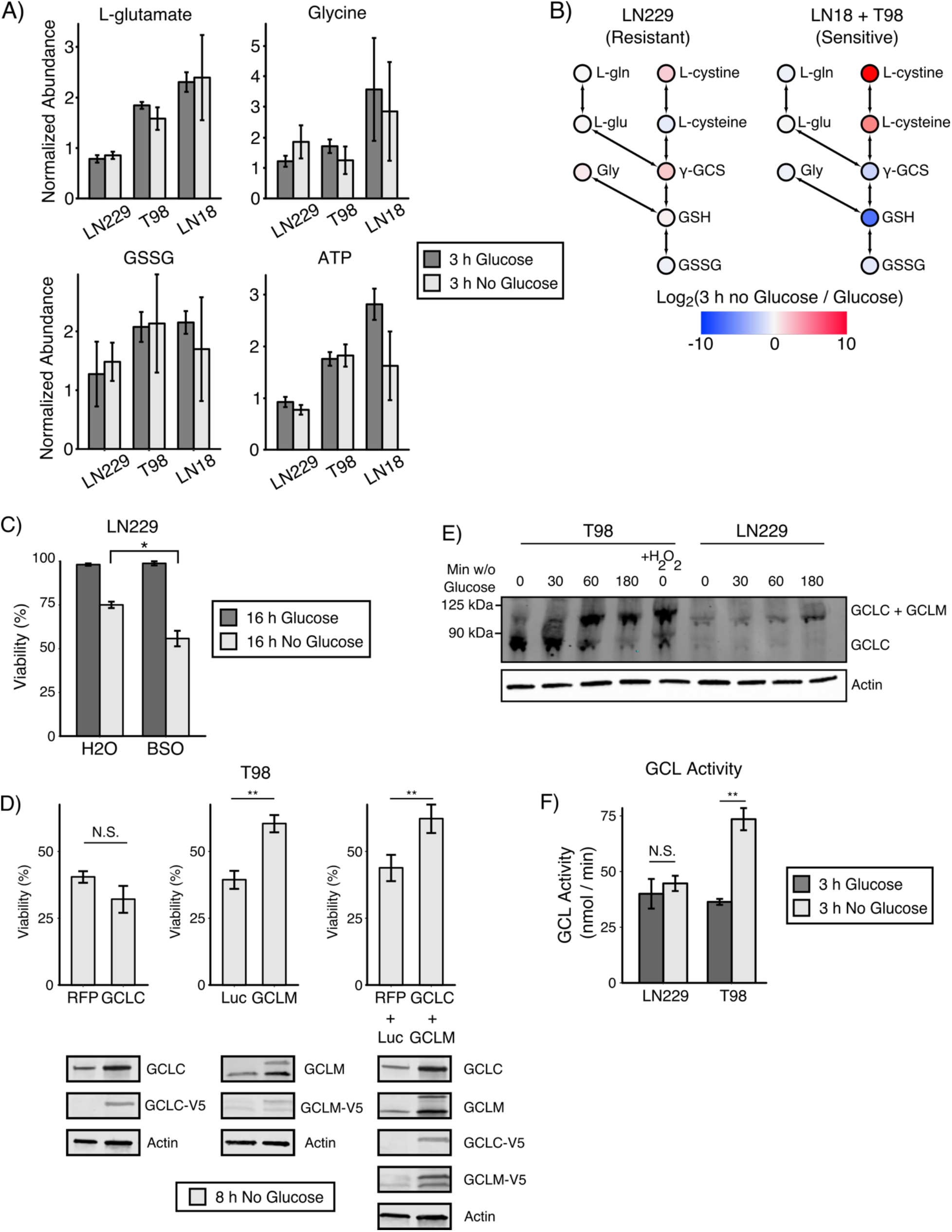
Glutamate-cysteine ligase (GCL) activity moderately regulates resistance to glucose deprivation. **A)** Individual bar plots from LC-MS metabolomics showing that the GSH synthesis precursors glycine and L-glutamate, as well as oxidized glutathione (GSSG) and ATP, were not significantly regulated by 3 h of glucose deprivation in either resistant (LN229) or sensitive cells (T98, LN18). **B)** Log_2_ fold change in metabolite pool sizes upon 3 h of glucose deprivation overlaid on the GSH synthesis pathway. Red and blue represent accumulation and depletion, respectively, as shown on the indicated color scale. Data from LN18 and T98 cells were averaged. L-gln, L-glutamine; L-glu, L-glutamate; Gly, glycine. **C)** Treatment with the GCL inhibitor buthionine sulfoximine (BSO, 500 µM) sensitized glucose deprivation-resistant LN229 GBM cells to glucose deprivation. Cells were treated with either solvent (water) or BSO in the presence or absence of glucose for 16 h, and viability was measured by trypan blue exclusion. * denotes p < 0.05 by Student’s t-test (n=2). **D)** Overexpression of GCLM, but not GCLC, confers resistance to glucose deprivation. Lentiviral vectors (pLX302 and pLX304) were used to overexpress either subunit of GCL. Glucose deprivation-sensitive T98 cells were starved of glucose for 8 h and viability was measured by trypan blue exclusion. Overexpression of GCLM, but not GCLC, increased survival. Combined overexpression of GCLC and GCLM did not confer additional resistance compared to GCLM alone. ** denotes p < 0.01 by Student’s t-test (n=3). **E)** The GCL holoenzyme is formed rapidly after glucose deprivation in both sensitive and resistant cells. The glucose deprivation-sensitive T98 and -resistant LN229 cell lines were starved of glucose for the indicated time and then lysed in a non-reducing lysis buffer. Treatment with 10 mM H2O2 for 10 min was included as a positive control. Cell lysates were run in native PAGE conditions, and formation of the GCL holoenzyme was monitored by formation of the GCLC:GCLM complex (101 kDa) using a GCLC antibody. **F)** GCL activity increases in response to glucose deprivation in sensitive T98 but not resistant LN229 cells. T98 and LN229 cells were starved of glucose for 3 h, lysates were collected, and GCL activity was measured using a fluorescence-based microtiter plate assay^38^. ** denotes p < 0.01 by Student’s t-test (n=2)

The ligation of L-glutamate and L-cysteine is catalyzed by glutamate-cysteine ligase (GCL), which consists of a catalytic subunit (GCLC) and a modifier subunit (GCLM). GCLM binding can increase the activity of GCL up to 10 fold^32^ by decreasing the Km for glutamate and increasing the Ki for GSH^33^. We thus first tested whether modulating GCL activity could affect sensitivity to glucose deprivation. Indeed, the GCL inhibitor buthionine sulfoximine (BSO) sensitized both resistant LN229 and HT161 cells to glucose deprivation (Fig. 2C and Supporting Fig. 2B). In addition, overexpression of GCLM, but not GCLC, using CRISPRa (dCas9-VPR^34^) promoted resistance in glucose deprivation-sensitive T98 and LN18 cells (Supporting Fig. 2C). Because the magnitude of GCLC and GCLM overexpression using CRISPRa was modest, we validated our CRISPRa findings with lentiviral overexpression of GCL subunits which we hypothesized would provide more substantial overexpression and thus perhaps exert a greater effect on glucose deprivation sensitivity. However, similar to our CRISPRa results, exogenous overexpression of GCLM but not GCLC conferred only a modest but significant increase in resistance to glucose deprivation in T98 cells (Fig. 2D). Notably, the increased GCLM overexpression achieved by lentiviral vectors relative to CRISPRa was not accompanied by increased resistance to glucose deprivation, suggesting that these cells have reached an upper limit to which GCLM can contribute resistance. To test whether GCLC had become limiting upon GCLM overexpression, we co-overexpressed GCLC and GCLM and did not observe increased resistance to glucose deprivation relative to GCLM alone (Fig. 2D). Taken together, this data suggests that GCL only modestly regulates sensitivity to glucose deprivation.

### Glutamate-cysteine ligase (GCL) activity is activated by glucose withdrawal

We next tested whether glucose deprivation regulates GCL activity. First, we examined whether glucose deprivation affected the expression of GCLC and GCLM. Western blotting revealed that neither GCLC nor GCLM expression was significantly changed following glucose deprivation (Supporting Fig. 2D). Interestingly, the sensitive cell lines T98 and U87 exhibited substantially higher GCLC and GCLM expression than the resistant cell line LN229. Next, using native PAGE gels to monitor the formation of the GCL holoenzyme, we found that 60 min of glucose deprivation increased holoenzyme formation in T98 sensitive cells to an extent similar to the positive control H_2_O_2_ (Fig. 2E)^35^. The resistant cell line LN229 also demonstrated formation of the GCL holoenzyme following glucose deprivation, although holoenzyme formation was slower (180 min) and substantially less expressed than in T98 sensitive cells. Finally, using an *in vitro* assay^36^, we directly tested GCL activity and found that glucose deprivation increased GCL activity in glucose deprivation-sensitive T98 but not -resistant LN229 cells (Fig. 2F). Although our metabolomic data had suggested GCL activity was inhibited in sensitive cells following glucose deprivation (Fig. 2B), these data demonstrate that GCL activity is in fact *<u>increased</u>* following glucose deprivation in sensitive cells. This, in turn, suggests the possibility that GSH depletion following glucose deprivation in sensitive cells occurs not because of inhibited GSH synthesis, but rather because of extremely rapid GSH consumption.

### NADPH is the limiting reducing agent in glucose-deprived cancer cells

In sensitive cells, it has been shown that glucose deprivation induces high levels of reactive oxygen species (ROS), and that elevated ROS levels are responsible for glucose deprivation-induced cell death^9, 23, 24^. We thus measured ROS levels at multiple time points following glucose deprivation. In sensitive T98 cells, ROS levels were increased as early as 10 min after glucose deprivation and steadily increased over two h (Fig. 3A and Supporting Fig. 3A). We next performed metabolomic profiling of sensitive T98 cells at the same time points (Supporting Table 4). Interestingly, we found that L-cystine and L-cysteine accumulated 8- and 6-fold, respectively, within only 10 min of glucose deprivation (Fig. 3B). In contrast, levels of GSH did not decrease until after 60 min of glucose deprivation, and levels of GSSG increased slightly over time after glucose deprivation (Fig. 3C). This result is consistent with our finding that GCL is active upon glucose deprivation (Fig. 2). Notably, other metabolic destinations of L-cysteine including cystathionine, taurine, and hypotaurine did not dramatically increase upon glucose deprivation (Supporting Fig. 3B), suggesting that L-cysteine is not diverted to other metabolic pathways in glucose-deprived cells. Taken together, these data demonstrate that intracellular accumulation of L-cystine, L-cysteine, and ROS precedes GSH depletion.

**Figure 3.**
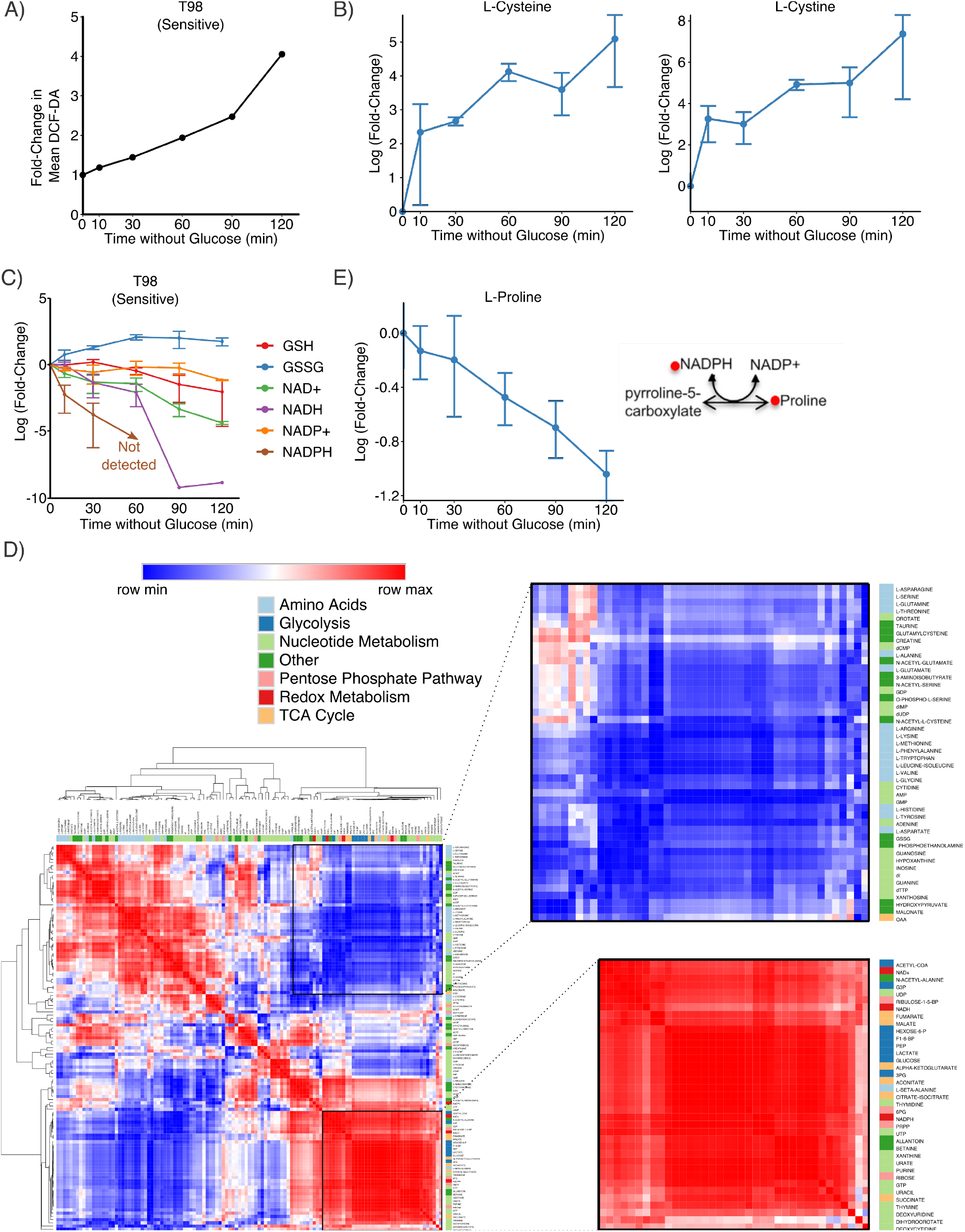
NADPH is the limiting reducing agent upon glucose deprivation. **A)** Reactive oxygen species (ROS) accumulate upon glucose deprivation. ROS was measured by flow cytometry using DCF-DA at the indicated time points. ROS accumulation steadily increased with time after glucose deprivation. **B)** Glucose deprivation-sensitive T98 cells were deprived of glucose and metabolites were extracted at the indicated time points. Accumulation of L-cystine and L-cysteine occurred within 10 min after glucose deprivation and steadily increased over the course of 2 h. **C)** NADPH is rapidly consumed following glucose deprivation. An LC-MS metabolomics method designed to detect NAD(P)H^72^ was used at the same time points as 3B. Depletion of NADPH preceded depletion of NADH and GSH, with NADPH dropping below the lower limit of detection within 60 min. **D)** The Pearson correlation coefficient was calculated between each metabolite, and hierarchical clustering was performed on the Pearson correlation coefficients. Clustering revealed that L-cystine, L-cysteine, and L-proline behave differently from all other proteinogenic amino acids. Glycolysis metabolites and reducing agents clustered together, reflecting that they both deplete upon glucose deprivation. 18 of 20 proteinogenic amino acids also clustered together, reflecting that amino acid metabolism and glycolysis are uncoupled upon glucose deprivation. **E)** L-proline is steadily depleted after glucose deprivation in sensitive T98 cells. The *de novo* synthesis of L-proline requires NADPH to convert pyrroline-5-carboxylate to L-proline.

NADPH is primarily generated by three main cellular metabolic sources: the oxidative pentose phosphate pathway (oxPPP), malic enzyme, and serine-derived one carbon metabolism^37^. Since glucose fuels both the oxPPP and the TCA cycle, we hypothesized that NADPH may be limiting when cells are deprived of glucose. We thus profiled the metabolome at the same time points using an LC-MS metabolomics method designed to detect NAD(P)H^38^. We found that levels of NADPH decreased dramatically within 10 min of glucose deprivation and were undetectable after 60 min (Fig. 3C). Notably, this short time scale is similar to the time required for L-cystine and L-cysteine accumulation following glucose deprivation. In contrast, intracellular pools of GSH and NADH were not depleted until later time points (90-120 min). These results suggest that NADPH, rather than NADH or GSH, is the limiting reducing agent when cancer cells are deprived of glucose.

To better understand the global metabolic changes upon glucose deprivation, we calculated the Pearson correlation coefficient between all pairs of metabolites. Hierarchical clustering of these Pearson correlation coefficients revealed a cluster of metabolites from glycolysis, the TCA cycle, and redox metabolism which are all rapidly depleted following glucose deprivation (Fig. 3D). A second large cluster included 18 of 20 proteinogenic amino acids and these metabolites were largely unperturbed by glucose deprivation. The only proteinogenic amino acids not found in this cluster were L-cysteine, which clustered with L-cystine, and L-proline which clustered with glycolytic and redox metabolites. Examination of L-proline levels revealed that intracellular concentrations of this amino acid steadily dropped after glucose deprivation (Fig. 3E). This may occur because DMEM does not contain L-proline, forcing cells to rely on *de novo* synthesis, but glucose-starved cells cannot provide the NADPH required for conversion of pyrroline-5-carboxylate to L-proline. Together, these results demonstrate that NADPH is the limiting reducing agent in glucose deprived cancer cells.

### L-cystine import, but not L-cysteine import, induces ROS mediated cell death in glucose-deprived cancer cells

The DMEM in which GBM cells are cultured contains 200 µM L-cystine but 0 µM L-cysteine. Thus, the rapid (<10 min) accumulation of L-cystine and L-cysteine following glucose deprivation is likely driven by L-cystine import and subsequent reduction to two molecules of L-cysteine. Notably, the reduction of L-cystine to L-cysteine consumes one reducing equivalent of NADPH^39^. Because NADPH is depleted on the same rapid time scale as L-cystine and L-cysteine accumulation (∼10 min), we hypothesized that the intracellular conversion of L-cystine to L-cysteine might drive glucose deprivation-induced NADPH depletion, ROS generation, and subsequent cell death. Therefore, we first tested whether L-cystine deprivation could rescue cells from glucose deprivation. Indeed, sensitive T98 cells exhibited very high viability after 8 h of glucose deprivation in cystine-free media, and supplementation with L-cystine restored glucose deprivation-induced cell death (Fig. 4A). Because GBM cells express L-cysteine transporters^40^ including EAAT3, ASCT1, SNAT1, and SNAT2, we next tested whether supplementation with L-cysteine, as opposed to L-cystine, could promote cell death in glucose starved cells. Although extracellular L-cysteine was consumed by T98 cells (Supporting Fig. 4A), L-cysteine supplementation did not increase cell death relative to cystine-free media. Notably, supplementation with L-cysteine did not confer greater resistance to glucose deprivation relative to L-cystine deprivation (data not shown). This may be due to the fact that L-cysteine spontaneously oxidized to L-cystine in cell culture media (not shown), suggesting that L-cysteine supplementation can result in L-cystine import. Thus, we tested whether supplementation with the GSH precursor downstream of L-cysteine, γ-GCS, could affect cell viability following glucose deprivation. Indeed, supplementation with γ-GCS in L-cystine-free media conferred a modest but significant resistance to glucose deprivation (Fig. 4B). These results demonstrate that L-cystine import, but not L-cysteine import, contributes to glucose deprivation-induced cell death.

**Figure 4.**
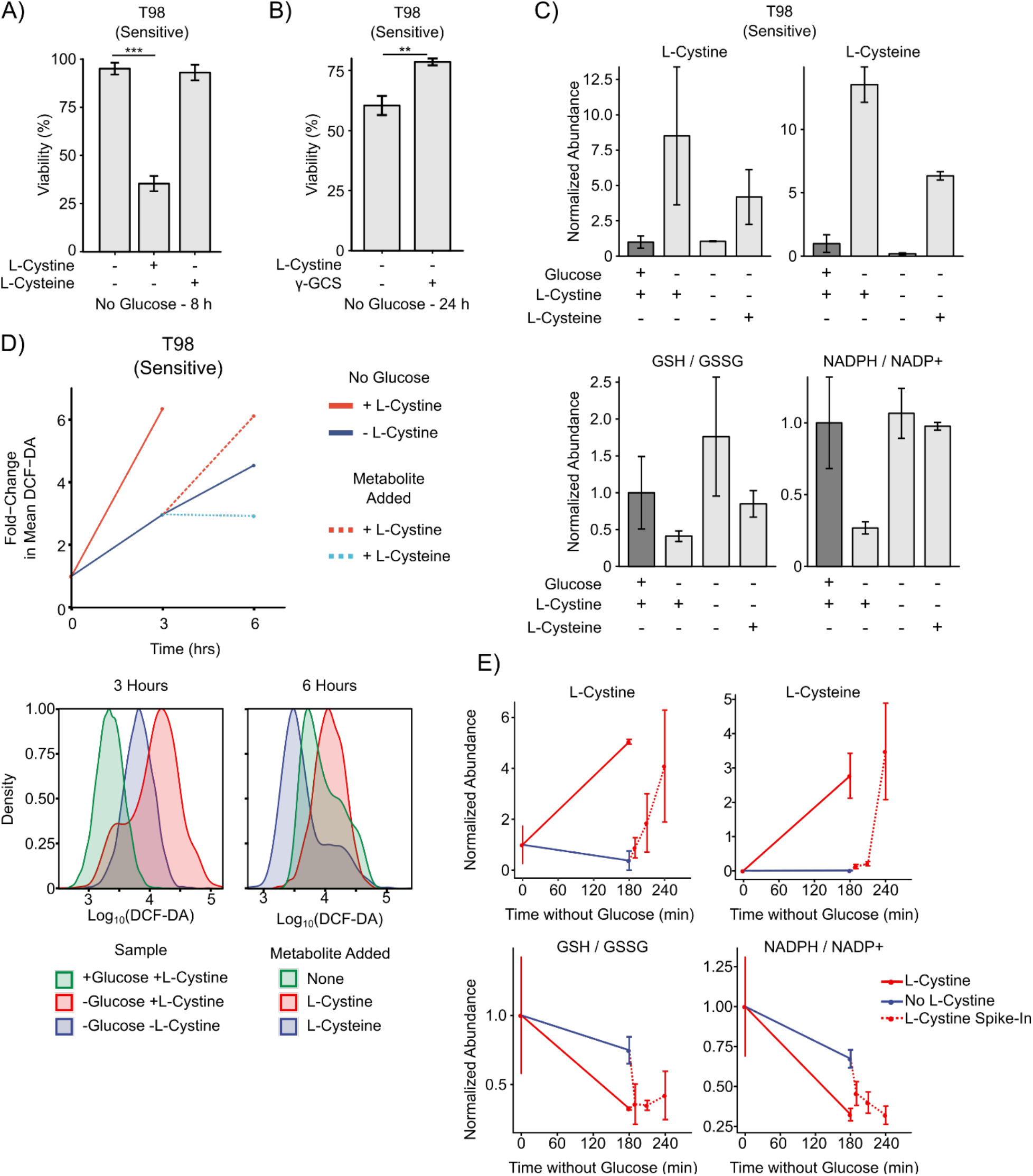
L-cystine import induces oxidative stress and ROS-mediated cell death in glucose deprived cancer cells. **A)** L-cystine, but not L-cysteine, import results in glucose deprivation-induced cell death. Glucose deprivation-sensitive T98 cells were cultured in L-cystine-free medium and subjected to glucose deprivation for 8 h. L-cystine starvation rescued T98 cells from glucose deprivation-induced cell death. Addition of L-cystine (200 µM), but not L-cysteine (200 µM), sensitized T98 cells to glucose deprivation. **B)** γ-GCS contributes resistance to glucose and L-cystine deprivation. Glucose deprivation-sensitive T98 cells were starved of glucose and L-cystine for 24 h. Addition of 200 µM γ-GCS conferred a modest resistance compared to DMSO control. ** denotes p-value < 0.01 by Student’s t-test (n = 3). **C)** L-cystine induces oxidative stress upon glucose deprivation. T98 cells were deprived of glucose in L-cystine-free medium for 3 h and metabolites were quantified using LC-MS metabolomics. Addition of L-cystine (200 µM), but not L-cysteine (200 µM), induced GSH depletion and oxidative stress as measured by the ratio of GSH / GSSG and NADPH / NADP+. **D)** ROS accumulation following glucose deprivation is driven by L-cystine import. T98 sensitive cells were deprived of glucose in the presence or absence of L-cystine for 3 h, and ROS levels were measured by flow cytometry by DCF-DA staining. Cells that had been starved of glucose and L-cystine for 3 h were then re-supplemented with L-cystine (200 µM), L-cysteine (200 µM), or neither for an additional 3 h. (Left) Mean fluorescent intensity of DCF-DA signal. (Center) Histograms of DCF-DA for T98 cells cultured with glucose, without glucose but with L-cystine, or without glucose or L-cystine for 3 h. (Right) Histograms of DCF-DA intensity for T98 cells starved of glucose and L-cystine for 3 h and then re-supplemented with L-cystine, L-cysteine, or neither for an additional 3 h. **E)** L-cystine import induces redox imbalance upon glucose deprivation. Glucose deprivation-sensitive T98 cells were starved of glucose for 3 h in the presence and absence of L-cystine, at which point L-cystine (200 µM) was spiked-in for 10, 30, and 60 min. Addition of L-cystine induced a redox imbalance as indicated by dramatic decreases in the ratios GSH / GSSG and NADPH / NADP+.

We next sought to understand the metabolic consequences of L-cystine and L-cysteine import upon glucose deprivation. Using LC-MS metabolomics, we found that L-cystine starvation prevented the accumulation of L-cystine and L-cysteine that normally occurs upon glucose deprivation in sensitive T98 cells (Fig. 4C). Supplementation of glucose-free media with L-cysteine instead of L-cystine led to increased intracellular levels of both L-cysteine and L-cystine, which may reflect either intracellular or extracellular oxidation of L-cysteine. We next asked whether the presence of L-cysteine or L-cystine during glucose starvation induced oxidative stress. Plotting the ratios of GSH/GSSG and NADPH/NADP+ indicated that sensitive T98 cells experienced oxidative stress (lower GSH/GSSG ratio and NADPH/NADP+ ratios) when deprived of glucose in media containing L-cystine but not in glucose-free media without L-cystine or glucose-free media supplemented with L-cysteine (Fig. 4C). Together, this indicates that the presence of L-cystine but not L-cysteine during glucose deprivation promotes GSH depletion and redox imbalance.

We next tested whether L-cystine import during glucose deprivation induces ROS. In glucose deprivation-sensitive T98 cells, 3 h of glucose deprivation in the presence of L-cystine induced a six-fold induction in ROS (Fig. 4D). Meanwhile, cells deprived of glucose in L-cystine-free medium exhibited only a three-fold induction of ROS, demonstrating that L-cystine import significantly contributes to ROS accumulation. We next sought to uncouple glucose deprivation from L-cystine import by starving cells of glucose and L-cystine for 3 h and then supplementing cells with either L-cystine or L-cysteine for an additional 3 h. Consistent with our hypothesis that L-cystine import drives ROS accumulation in glucose-starved cells, L-cystine supplementation increased ROS levels. Addition of L-cysteine, in contrast, decreased ROS levels compared to an additional 3 h without L-cystine or glucose. Next, we used LC-MS metabolomics to profile the metabolic changes that occurred after supplementation of L-cystine. Again, sensitive T98 cells were starved of glucose and L-cystine for 3 h, and then L-cystine was spiked into the cell culture media. Following L-cystine addition, levels of intracellular L-cystine and L-cysteine levels accumulated rapidly, with levels increasing within just 10 min of L-cystine spike-in (Fig. 4E). In addition, the ratios of GSH/GSSG and NADPH/NADP^+^ were dramatically reduced within 10 min of L-cystine spike-in. Taken together, these data demonstrate that import of L-cystine, but not L-cysteine, promotes ROS accumulation, GSH depletion, and redox imbalance in sensitive cancer cells deprived of glucose.

### L-cystine import from the glutamate/cystine antiporter xCT/SLC7A11 promotes cell death following glucose starvation

The primary source for intracellular L-cystine is the glutamate/cystine antiporter, system x_c_^-^. System x_c_^-^ comprises a heterodimer containing a non-specific heavy chain, SLC3A2, and a specific light chain, xCT/SLC7A11. Recent reports have shown that system x_c_^-^ activity promotes cancer cell dependency on glucose^14, 16, 24^, although the mechanism by which system x_c_^-^ promotes glucose deprivation-induced cell death remains disputed. Therefore, we sought to test the role of system x_c_^-^ in L-cystine accumulation and cell death observed following glucose deprivation. First, by Western blotting, we found that expression of xCT/SLC7A11 correlated with sensitivity to glucose deprivation in both glioblastoma and sarcoma cell lines (Fig. 5A and Supporting Fig. 5A). Next, we tested whether pharmacological and genetic inhibition of system x_c_^-^ affected sensitivity to glucose deprivation. Indeed, pharmacological inhibition of xCT with either sulfasalazine (SASP) or erastin prevented glucose deprivation-induced cell death (Fig. 5B-C and Supporting Fig. 5B). In addition, knockdown of xCT with CRISPRi^41^ (dCas9-KRAB) rescued LN18 cells from glucose deprivation (Fig. 5D). These findings corroborate recent reports that system x_c_^-^ activity and expression is required for glucose deprivation-induced cell death^14, 16, 24^.

**Figure 5.**
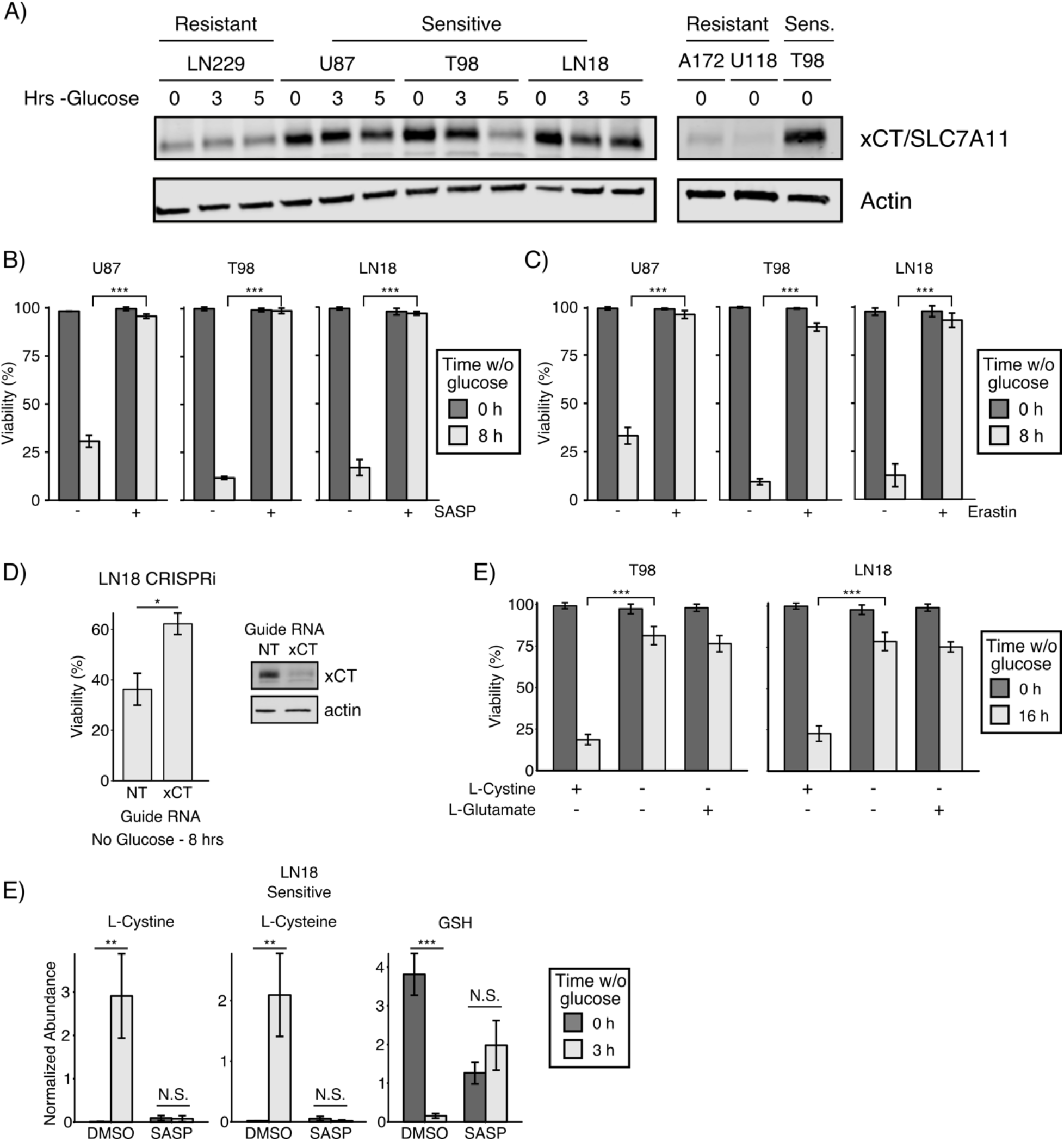
L-cystine import from the glutamate/cystine antiporter, system xc-, is required for glucose deprivation-induced cell death. **A)** Expression of the specific light chain of system x_c_^-^, xCT/SLC7A11, correlates with sensitivity to glucose deprivation in GBM cell lines. The resistant and sensitive GBM cell lines were starved of glucose for the indicated times and then lysed. Expression of xCT/SLC7A11 was assessed by Western blotting. Actin was used as an equal loading control. **B)** Pharmacological inhibition of xCT rescued sensitive cells from glucose deprivation-induced cell death. The indicated sensitive GBM cells were treated with either sulfasalazine (SASP, 500 µM) or erastin (10 µM) in the presence or absence of glucose for 8 h, and viability was assessed by trypan blue exclusion. *** denotes p < 0.001 by Student’s t-test (n=3). **C)** Genetic knockdown of xCT promoted resistance to glucose deprivation in sensitive LN18 cells. Cells were first infected with CRISPRi machinery (dCas9-KRAB^42^) and then infected with a second vector carrying the indicated guide RNA. Cells were then cultured 8 h in the presence or absence of glucose, and viability was measured by trypan blue exclusion. * denotes p < 0.05 by Student’s t-test (n=3). Western blotting with antibodies against xCT and the equal loading control actin confirmed knockdown. NT, non-targeting control. **D)** L-cystine starvation rescues sensitive cells from glucose deprivation-induced death. The sensitive GBM cell lines were cultured in L-cystine-free DMEM in the presence and absence of glucose for 16 h. The media was supplemented with either L-cystine (200 µM) or L-glutamate (100 µM), and viability was measured by trypan blue exclusion. *** denotes p < 0.001 by Student’s t-test (n=3). **E)** Inhibition of the xCT cystine/glutamate antiporter prevents GSH depletion following glucose deprivation. The sensitive GBM cell line LN18 was cultured with and without glucose in the presence or absence of the xCT inhibitor sulfasalazine (SASP, 500 µM) for 3 h, and then intracellular metabolite concentrations were measured using LC-MS metabolomics. xCT inhibition prevented the accumulation of L-cystine and L-cysteine and depletion of GSH following glucose deprivation. ** denotes p < 0.01, *** denotes p < 0.001 by Student’s t-test (n=3).

Because system x_c_^-^ both imports L-cystine and exports L-glutamate, we next tested whether L-glutamate export, in addition to L-cystine import, plays a role in glucose deprivation-induced cancer cell death. We thus cultured cells in media with L-cystine but without L-glutamate, media lacking L-cystine and L-glutamate, or media lacking L-cystine but supplemented with L-glutamate (i.e., mimicking system x_c_^-^ activity). In both T98 and LN18 sensitive GBM cells, L-cystine starvation rescued cells from glucose deprivation-induced cell death, but L-glutamate supplementation did not restore cell death (Fig. 5E). These results demonstrate that import of L-cystine from system x_c_^-^, but not export of L-glutamate, regulates glucose deprivation-induced cell death.

To further understand the metabolic effect of system x_c_^-^ activity upon glucose deprivation, we performed LC-MS metabolomics on sensitive LN18 cells deprived of glucose in the presence or absence of the xCT inhibitor SASP. As expected, SASP treatment prevented the accumulation of L-cystine and L-cysteine following glucose deprivation (Fig. 5F). In the presence of SASP, the basal concentration of GSH in cells cultured with glucose was reduced 67%, reflecting the fact that SASP-treated cells must rely on *de novo* synthesis of L-cysteine (rather than L-cystine import) in order to synthesize GSH. Surprisingly, when cells were starved of glucose in the presence of SASP, the levels of GSH increased rather than decreased (Fig. 5F). Taken together with our metabolomic profiling of L-cystine starved cells (Fig. 4C), these results demonstrate that system x_c_^-^ activity promotes glucose deprivation-induced cell death through import of L-cystine, depletion of GSH and consumption of NADPH by L-cystine reduction to L-cysteine.

### Inhibition of glucose metabolism is synthetically lethal with inhibition of GCL

Building upon our observations that depletion of NADPH and GSH promotes glucose deprivation-induced cell death, we sought to identify drug combinations that mimic this phenotype in the presence of glucose. To identify candidate glycolytic nodes to target, we analyzed data from the Cancer Dependency Map (DepMap^42^) to identify glycolytic genes whose CERES dependency score was correlated with high xCT expression in glioblastoma (GBM) and Ewing’s sarcoma cell lines, since xCT expression correlates with sensitivity to glucose deprivation in those cell types (Fig. 5A and Supporting Fig. 5A). Of 22 glycolytic enzymes, we found that xCT expression was most strongly correlated with dependency on the glucose transporter GLUT1 (*SLC2A1*) in GBM and also strongly correlated with dependency on GLUT1 in sarcoma cell lines (i.e., high xCT expressing cells have large negative CERES scores, indicating greater dependency on GLUT1) (Fig. 6A and Supporting Fig. 6A-C). We thus hypothesized that glucose deprivation sensitive GBM and sarcoma cell lines would be sensitive to combined inhibition of GLUT1 and GSH synthesis. To test this hypothesis, we treated T98 sensitive GBM cells with increasing concentrations of an inhibitor of GLUT1 (STF-31^18^), an inhibitor of GCL (BSO), or the combination of both inhibitors. Treatment with either drug alone showed little effect on viability (Fig. 6B), whereas co-inhibition of GLUT1 and GCL synergistically induced cell death in T98 cells, as measured by calculating the Combination Index^43^ (CI < 1 indicates synergy). Because STF-31 has been reported to inhibit both GLUT1^18^ and NAMPT^44^, we investigated whether NAMPT may play a role in combined STF-31 and BSO treatment. Using the DepMap data, we asked whether NAMPT dependency correlated with xCT expression but found no correlation between NAMPT dependency and xCT expression in either GBM or Ewing’s sarcoma cell lines (Supporting Fig. 6D). Taken together, these results demonstrate that combined inhibition of GLUT1 and GCL provoke synergistic cell death in T98 cells.

**Figure 6.**
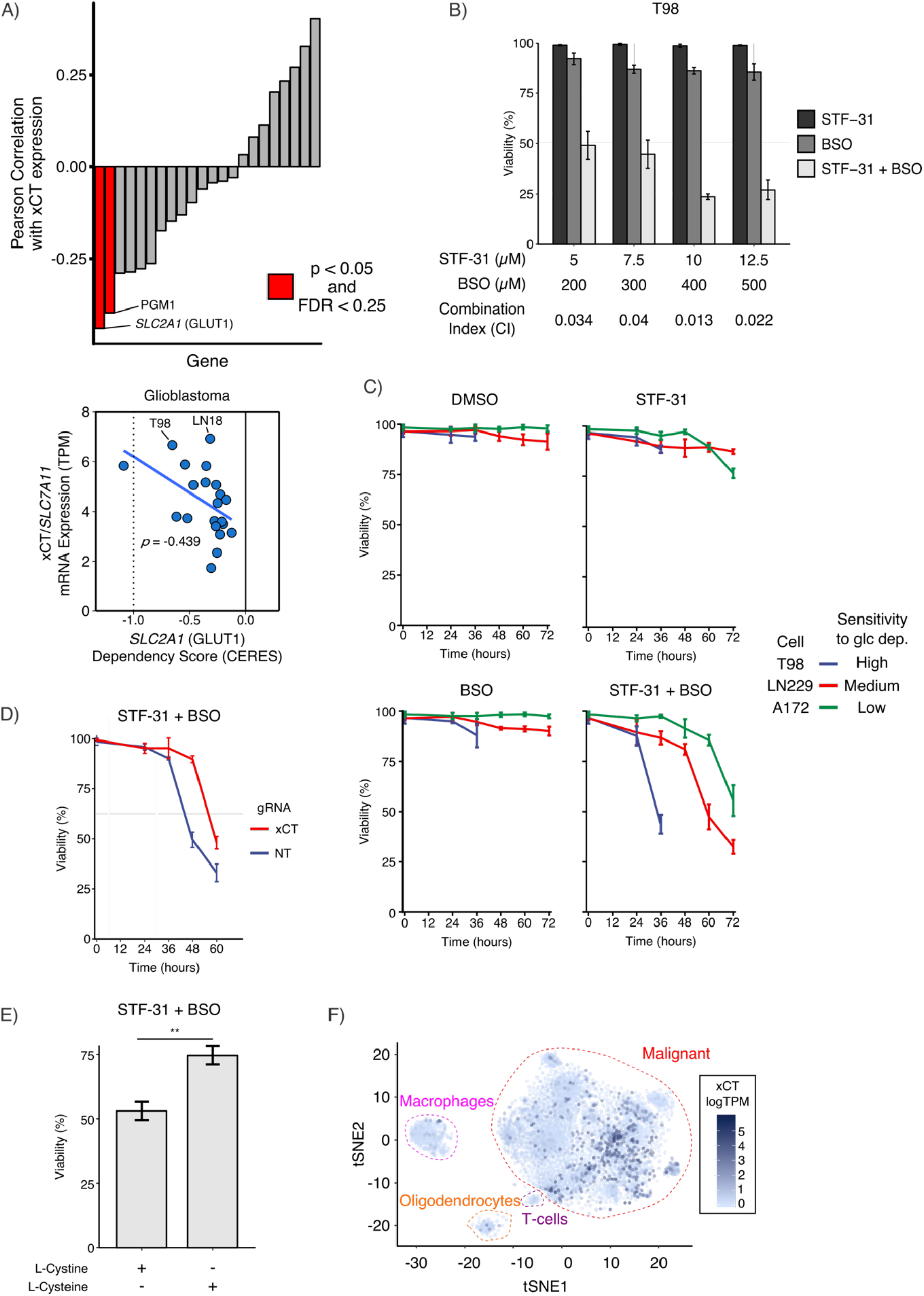
Inhibition of glutathione synthesis is synthetically lethal with GLUT1 inhibition. **A)** Glioblastoma cells rely on SLC2A1 (GLUT1) as xCT expression increases. Using data from the Cancer Dependency Map (DepMap^43^), glycolytic genes were filtered for expression in GBM and then ranked by their Pearson correlation coefficient between dependency (CERES score) and xCT expression. A negative CERES score indicates increased dependency. *SLC2A1* (GLUT1) and PGM1 were the top hits for GBM. **B)** Co-inhibition of GLUT1 and GCL synergistically induces cell death in glucose deprivation-sensitive cells. T98 cells were cultured in 5 mM glucose and treated with STF-31 and/or BSO at the indicated doses. After 36 h of treatment, the combination of drugs synergistically induced cell death. Combination Index values < 1 indicate synergy^72^. **C)** Sensitivity to glucose deprivation correlates with response to combined STF-31 and BSO treatment. Highly glucose deprivation-sensitive T98, medium-resistant LN229, and highly-resistant A172 GBM cells were cultured in 5 mM glucose and treated with STF-31 (12.5 µM), BSO (500 µM), or both. T98 cells died after 36 h of treatment, whereas LN229 cells died after 60 h of treatment, and A172 cells died after 72 h of combined STF-31 and BSO treatment. **D)** Genetic knockdown of xCT confers resistance to STF-31 and BSO treatment in xCT-high LN18 cells. Cells were first infected with CRISPRi machinery (dCas9-KRAB^45^) and then infected with a second vector carrying the indicated guide RNA. Cells were cultured in 5 mM glucose and treated with STF-31 (12.5 µM), BSO (500 µM), or both. After 48 h, combined STF-31 and BSO treatment induced cell death in non-targeting control cells but not in xCT-knockdown cells. **E)** L-cystine import contributes to combined STF-31 and BSO-induced cell death in T98 cells. Cells were cultured in 5 mM glucose supplemented with either L-cystine (200 µM) or L-cysteine (200 µM). L-cysteine treatment conferred resistance to combined STF-31 (12.5 µM) and BSO (500 µM) treatment. ** denotes p < 0.01 by Student’s t-test (n=3). **F)** xCT expression is increased in a subset of GBM. Single-cell RNA-sequencing data from 28 adult and pediatric patients^45^ was analyzed for xCT expression (logTPM). xCT expression was overlaid onto a tSNE plot to compare malignant GBM cells to macrophages, T cells, and oligodendrocytes

We next sought to test whether sensitivity to glucose deprivation correlated with response to combined GLUT1 and GCL inhibition. Using a panel of GBM cell lines with either high (T98), medium (LN229), or low (A172) sensitivity to glucose deprivation, we treated cells with STF-31, BSO, or the combination of both drugs. Treatment with individual drugs did not cause significant cell death in any of the three cell types. However, at 36 h, highly glucose deprivation sensitive T98 cells treated with the combination of STF-31 and BSO exhibited a dramatic reduction in cell viability compared to either treatment alone (Fig. 6C). Medium glucose deprivation sensitive LN229 cells also responded to the drug combination but only after 60 h of treatment. Finally, low glucose deprivation sensitivity A172 cells showed a modest response to combined STF-31 and BSO treatment at 72 h. Similar results were observed in sarcoma cell lines with high and medium sensitivity to glucose deprivation (Supporting Fig. 6E). Together, these results demonstrate that sensitivity to glucose deprivation correlates with sensitivity to a drug combination blocking glucose transport (GLUT1) and GSH synthesis (GCL).

Next, to test the role of system x_c_^-^ in response to combined GLUT1 and GCL inhibitors, we evaluated the response to the drug combination in xCT knockdown cells. In sensitive LN18 cells, we found that xCT knockdown conferred resistance to combined STF-31 and BSO treatment compared to non-targeting control (Fig. 6D). We next tested whether L-cystine import, and subsequent NADPH consumption, contributed to cell death in combined STF-31 and BSO treatment. Culturing T98 cells with L-cysteine rather than L-cystine again conferred resistance to the drug combination (Fig. 6E), suggesting that L-cystine reduction contributes to cell death upon combined STF-31 and BSO treatment. These data further support that L-cystine import from xCT is important for response to combined GLUT1 and GCL inhibition.

To further evaluate whether combined STF-31 and BSO treatment mimics glucose deprivation-induced cell death, we measured ROS accumulation in cells treated with the drug combination. As expected, 24 h of BSO treatment induced a 3-fold increase in ROS accumulation, reflecting the fact that GSH synthesis is inhibited (Supporting Fig. 6F). In contrast, 24 h of STF-31 treatment alone did not induce ROS accumulation. However, combined STF-31 and BSO treatment induced a 4-fold increase in ROS accumulation. Taken together, these data suggest that combined inhibition of GLUT1 and GCL mimics glucose deprivation-induced cell death.

We next investigated xCT expression within GBM tumors using single-cell RNA-sequencing data from 28 adult and pediatric patients^45^. Malignant GBM cells exhibited higher xCT expression when compared to macrophages, T cells, and oligodendrocytes (Fig. 6F). Examining xCT expression in individual patients revealed that malignant GBM cells exhibited significantly higher expression of xCT than non-malignant cells in 10 out of 20 patients. Together, this illustrates the potential of xCT as a therapeutic target for a subset of GBM patients (Supporting Fig. 7).

Having validated that combined inhibition of GLUT1 and GCL synergistically induced cell death in xCT-high GBM and Ewing’s sarcoma cell lines, we next sought to identify other cancer types which might exhibit sensitivity to this drug combination. Mining the DepMap database, we found that melanoma and lung adenocarcinoma cancer cell lines exhibited a strong negative correlation between GLUT1 dependency and xCT expression (Supporting Fig. 8A), suggesting that these cells lines would respond synergistically to GLUT1 and GCL co-inhibition. In contrast, high xCT expressing cell lines derived from breast, kidney, ovarian, and colorectal cancers did not exhibit increased dependence on GLUT1, suggesting that high xCT expressing cells from these tumor types would not synergistically respond to combined GCL and GLUT1 inhibition. To test this hypothesis, we treated the breast cancer cell lines M453 (xCT low) and M436 (xCT high) with STF-31 and BSO. We found that neither breast cancer cell line exhibited significant cell death in response to combined treatment even at 72 h (Supporting Fig. 8B). Taken together, these data demonstrate the ability of the DepMap data to predict response to co-inhibition of GLUT1 and GCL.

## Discussion

The metabolic reprogramming that occurs during oncogenesis can result in increased dependency on certain substrates for survival (e.g., glucose and glutamine)^7, 46^. As such, therapeutic strategies that target altered glucose metabolism in cancer cells have attracted considerable attention^1–5, 47^. Here, we demonstrate a synthetic lethal therapeutic strategy targeting the interconnected nature of glucose and redox metabolism. Our results show that glucose deprivation-induced cell death is driven not by lack of glucose, but rather by NADPH depletion following L-cystine import and reduction to L-cysteine. Consistent with this mechanism, glucose deprivation-induced cell death requires activity of the L-glutamate/L-cystine antiporter xCT. By mining the cancer dependency database DepMap, we found that GBM cells with high xCT expression are sensitive to depletion of the glucose transporter GLUT1. Building on these observations, we demonstrate that co-targeting GLUT1 and GSH metabolism induced a synthetic lethal interaction in glucose deprivation-sensitive glioblastoma and sarcoma cell lines. Taken together, our results demonstrate a synthetic lethal approach to exploit the glucose addiction phenotype in cancer cells.

Our results add to the growing literature surrounding synthetic lethal targeting of tumor metabolism. For example, passenger genomic deletions of metabolic enzymes, including enolase 1^48^, malic enzyme 2^49^, or methylthioadenosine phosphorylase (MTAP)^50–52^, can create synthetic lethal vulnerabilities unique to cancer cells. Similarly, in B cell malignancies, transcriptional repression of the pentose phosphate pathway creates a cancer cell-specific lack of antioxidant protection^53^, whereas in renal cell carcinomas, VHL mutation creates a sensitivity to inhibition of the glucose transporter GLUT1^54^. In addition, our work builds upon previous studies demonstrating that co-targeting glucose metabolism and redox metabolism can induce cell death in some cellular contexts. For example, the glycolytic inhibitor 2-deoxyglucose (2-DG) demonstrates enhanced cytotoxicity when paired with BSO in human breast cancer cells^55^ or with antimycin A in human colon cancer cells^20^. Interestingly, however, 2-DG supplementation can rescue cancer cells in the absence of glucose^9, 56^. This may be due to the fact that 2-DG can be metabolized in the pentose phosphate pathway^57^ (PPP), thus allowing 2-DG to serve as an alternative source of NADPH when glucose is restricted. Our results provide a rationale for co-targeting GLUT1 and GSH metabolism in high xCT expressing GBM and Ewing’s sarcomas.

Multiple recent studies have demonstrated that the L-glutamate/L-cystine antiporter xCT is required for cell death upon glucose withdrawal^14, 16, 24^, although the role of xCT is debated. Because xCT exports L-glutamate and imports L-cystine, studies have suggested that xCT promotes glucose addiction by depleting L-glutamate and thereby impairing the TCA cycle^14, 16^. However, in our GBM cell lines, we find no evidence of TCA cycle impairment following glucose starvation. Notably, in glucose withdrawal-sensitive cells, the forward flux through the TCA cycle was increased and levels of L-glutamate were unchanged following glucose deprivation (Figs. 1D and 2A). In contrast, our data suggest that xCT promotes L-cystine import and subsequent depletion of NADPH as L-cystine is reduced to L-cysteine, consistent with the model proposed by Goji et al.^24^. Notably, the depletion of NADPH by L-cystine import is consistent with our observation of increased flux through the TCA cycle following glucose deprivation, as cells attempt to generate NADPH through malic enzyme. This finding is consistent with a recent report that knockdown of malic enzyme can sensitize cancer cells to glucose deprivation^58^. Finally, although it has been shown that glucose deprivation induces oxidative stress in sensitive cells^23, 59–61^, the source of reactive oxygen species (ROS) has remained unclear. Our results suggest that L-cystine reduction is a primary source of redox imbalance, thereby allowing enabling accumulation of ROS from sources including mitochondria and NADPH oxidase^12^.

Notably, targeting GSH synthesis has long been viewed as a potential cancer therapy^62, 63^, and the GCL inhibitor BSO has been used in combination with the alkylating agent melphalan in clinical trials with mixed results (NCT00005835 and NCT00002730)^64, 65^. Our findings suggest that pairing BSO with inhibitors of GLUT1 may prove synthetically lethal in cancers with high xCT expression including glioblastomas, sarcomas, melanomas, and lung adenocarcinomas. Because high xCT levels are associated with poor prognosis in a number of cancer types, including GBM^66^ and triple-negative breast cancer^67^, the xCT transporter has been suggested as a therapeutic target for cancer^67–69^. Additionally, engineered enzymes that degrade extracellular L-cystine and L-cysteine and thereby indirectly inhibit xCT, blocking intracellular GSH production and upregulating ROS, have shown promise in preclinical models^70^. Our findings, in contrast, suggest that not all tumors may benefit from xCT inhibition, particularly tumor cells in nutrient-limited microenvironments. In fact, our results suggest that pairing xCT inhibitors with GLUT1 inhibitors would be counterproductive, leading to greater tumor cell survival. The L-cystine dependent mechanism described here, in addition to findings that GLUT1 is commonly upregulated in GBM^71^, suggests that a synthetic lethal approach targeting GSH synthesis and GLUT1 activity may result in selective toxicity towards cancer cells.

In summary, the highly interconnected nature of redox homeostasis and glycolysis in cancer cells provides an opportunity for synthetic lethal drug combinations targeting GSH synthesis and glucose metabolism. Future studies will be necessary to evaluate the efficacy and toxicity of combined GLUT1 and GCL inhibition *in vivo* and also to understand why some tumor types (e.g., GBM and Ewing’s sarcoma) but not others (e.g., breast cancer) are susceptible to this drug combination. Taken together, these results demonstrate the power of integrating metabolomic profiling with public databases of cancer dependencies (e.g., DepMap) to identify synthetic lethal vulnerabilities of cancer cells.

## Methods

### Cell Culture

All cell lines were cultured in high-glucose DMEM (4.5 g/l glucose, 110mM pyruvate; Mediatech) supplemented with 10% (v/v) FBS (Omega Scientific) plus 1% (v/v) anti-anti (Invitrogen). For glucose starvation, cells were washed twice with PBS and then incubated in DMEM without glucose and pyruvate (0 g/l glucose, 0 mM pyruvate; Invitrogen) supplemented with 10% dialyzed FBS (Omega Scientific) plus 1% anti-anti. GBM cell lines LN18, LN229, T98, and U87 and sarcoma cell lines HT161 and TC32 were a gift from Thomas Graeber (University of California, Los Angeles). GBM cell lines A172 and U118MG were gifts from David Changhan Lee (University of Southern California) and Matthew Lazzara (University of Virginia), respectively.

### Genetic modifications

Knockdown and overexpression of genes using CRISPR interference^41^ and CRISPR activation^34^ was done as previously described. Briefly, guide RNAs were expressed using pLenti-hygro-mTagBFP2 and paired with either lenti-EF1a-dCas9-KRAB-Puro or lenti-EF1a-dCas9-VPR-Puro. Wild-type GCLM (pLX304), GCLC (pLX302), luciferase (pLX304), or RFP (pLX302) was expressed using lentiviral vectors. Following infection, cells were selected using hygromycin (pLenti-hygro-mTagBFP2), puromycin (lenti-EF1a-dCas9-VPR-Puro or lenti-EF1a-dCas9-KRAB-Puro or pLX302), or blasticidin (pLX304). For CRISPRa experiments, guide RNAs were designed to target upstream of the transcriptional start site. For CRISPRi experiments, guide RNAs were designed to target downstream of the transcriptional start site.

### Cell Viability Analysis

Viability was measured by Trypan blue exclusion using a TC20 automated cell counter (BioRad) using optimized, cell type-specific imaging parameters to distinguish live and dead cells.

### Western blotting

Cells were lysed in modified RIPA buffer (50 mM Tris–HCl (pH 7.5), 150 mM NaCl, 10 mM β-glycerophosphate, 1% NP-40, 0.25% sodium deoxycholate, 10 mM sodium pyrophosphate, 30 mM sodium fluoride, 1 mM EDTA, 1 mM vanadate, 20 mg/ml aprotinin, 20 mg/ml leupeptin, and 1mM phenylmethylsulfonyl fluoride). Whole cell lysates were resolved by SDS–PAGE on 4–15% gradient gels and blotted onto nitrocellulose membranes (Bio-Rad). Native lysates were collected in a native sample buffer (50 mM tris, pH 8.0, 1% NP-40, 150 mM NaCl) containing protease and phosphatase inhibitors. Native PAGE gels were run in Tris/Glycine/SDS at 4C without boiling or addition of reducing agents. Membranes were blocked over-night and then incubated sequentially with primary and IRDye-conjugated secondary antibodies (Li-Cor). Blots were imaged using the Odyssey Infrared Imaging System (Li-Cor).

### Liquid Chromatography-Mass Spectrometry Metabolomics

Cell lines were plated onto 6-well plates at a density of 1 x 10^5^ cells/well. Cells were washed twice with PBS before being treated with media. Metabolite extraction was performed 3 h after adding glucose starvation unless otherwise specified. For extraction of intracellular metabolites, cells were washed on ice with 1 mL ice-cold 150 mM ammonium acetate (NH_4_AcO, pH 7.3). 1 mL of −80 °C 80% MeOH was added to the wells, samples were incubated at −80 °C for 20 min, then cells were scraped off and supernatants were transferred into microcentrifuge tubes. Samples were pelleted at 4C for 5 min at 15k rpm. Supernatants were transferred into LoBind Eppendorf microcentrifuge tubes and the cell pellets were re-extracted with 200 uL ice-cold 80% MeOH, spun down and the supernatants were combined. Metabolites were dried at room temperature under vacuum and re-suspended in water for injection. For extraction of extracellular metabolites, 20 uL of cell free-blank and conditioned media samples were collected from wells. Metabolites were extracted by adding 500 ul −80C 80% MeOH, dried at room temperature under vacuum and re-suspended in water for injection.

NAD(P)H metabolites were extracted using a method described here^38^. Briefly, for extraction of intracellular metabolites, a two solvent method is used. Solvent A is 40:40:20 acetonitrile:methanol:water with 0.1 M formic acid, and solvent B is 15% NH_4_HCO_3_ in water (w:v), precooled on ice. Solvent A was added to the cell pellet, vortexed for 10 s, and allowed to sit on ice for 3 min. For each 100 μl of solvent A, 8.7 μl of solvent B was then added and vortexed to neutralize the sample, and the mixture allowed to sit on dry ice for 20 min. Volumes were calculated to produce a total volume of 50 μl solvent per 1 μl of cell volume. Samples were then centrifuged at 16,000 *g* for 15 min at 4°C and supernatant taken for to be dried at room temperature under vacuum and resuspended in water for LC-MS analysis.

Samples were randomized and analyzed on a Q-Exactive Plus hybrid quadrupole-Orbitrap mass spectrometer coupled to an UltiMate 3000 UHPLC system (Thermo Scientific). The mass spectrometer was run in polarity switching mode (+3.00 kV/-2.25 kV) with an m/z window ranging from 65 to 975. Mobile phase A was 5 mM NH4AcO, pH 9.9, and mobile phase B was acetonitrile. Metabolites were separated on a Luna 3 μm NH2 100 Å (150 × 2.0 mm) column (Phenomenex). The flowrate was 300 μl/min, and the gradient was from 15% A to 95% A in 18 min, followed by an isocratic step for 9 min and re-equilibration for 7 min. All samples were run in biological triplicate.

Metabolites were detected and quantified as area under the curve based on retention time and accurate mass (≤ 5 ppm) using the TraceFinder 3.3 (Thermo Scientific) software. Raw data was corrected for naturally occurring 13C abundance (49). Extracellular data was normalized to integrated cell number, which was calculated based on cell counts at the start and end of the time course and an exponential growth equation. Intracellular data was normalized to the cell number and cell volume at the time of extraction. Pathway maps were made with Cytoscape software (50).

### Flow cytometry

Cells were incubated with either 5 mM CM-H2DCFDA for 10min before analysis using a MACSQuant® Analyzer 10 Flow Cytometer. Cells were gated using forward scatter and side scatter to remove debris and dead cells. To quantify changes in DCF-DA signal, mean fluorescent intensity after gating was used.

### GCL activity assay

Enzymatic activity for GCL was assessed using a naphthalene dicarboxaldehyde (NDA) derivatization method to form cyclized, fluorescent products^36^. Native lysates were collected and samples were equally loaded to total protein content. Samples were split into 2 samples (A and B). A GCL reaction cocktail (400 mM Tris, 40 mM ATP, 40 mM Glutamate, 40 mM Cysteine, 2 mM EDTA, 20 mM sodium borate, 2 mM serine, 40 mM MgCl2) was added to sample A. Lysis buffer was added to sample B for background measurements. Samples were incubated at 37C for 15 min, followed by quenching with 200 mM 5-sulfosalicyclic acid (SSA). Samples were placed on ice for 20 min then centrifuged at 2,000g for 10 min at 4C. The supernatant was collected and 20 µL was transferred to a 96 well plate designed for fluorescent detection. A standard curve of γ-GCS was prepared (0 to 140 µM). 20 µL of each standard solution was added to the 96 well plate. 180 µL of NDA derivatization buffer was added to each well, followed by incubation for 30 min at RT in the dark. NDA-γ-GCS was measured by a plate reader (472 nm excitation / 528 nm emission).

### Bioinformatic analysis

Data from the cancer dependency map (DepMap^42^) was downloaded from the depmap data portal (https://depmap.org/portal/). Cell lines were grouped by their cancer subtype and dependencies scores of genes from glycolysis were correlated against xCT expression. Significance was assessed by permutations shuffling dependency scores and xCT expression. Single-cell RNA-sequencing data from adult and pediatric glioblastoma tumors^45^ was downloaded from the Broad institute single cell portal (https://portals.broadinstitute.org/single_cell). Cell assignment labels were taken from the original publication.

## Acknowledgments

This work supported by grant #IRG-16-181-57 from the American Cancer Society, The Rose Hills Foundation, The USC Provost’s Office, and The Viterbi School of Engineering. J.H.J. was supported by a Mork Family Doctoral Fellowship. We would like to thank Melanie MacMullan and Pin Wang for assistance with flow cytometry experiments and Matt Pratt and Chao Zhang for scientific discussion.

## Author contributions

J.H.J. and N.A.G. conceived the project. J.H.J., P.S.P., S.P. conducted the experiments. A.D. aided with LC-MS metabolomics methodology. J.H.J. and N.A.G. interpreted data and wrote the manuscript.

