## Supporting Figures for "NADPH consumption by L-cystine reduction creates a metabolic vulnerability upon glucose deprivation"

Running title: *L-cystine drives glucose deprivation-induced cell death*

To whom correspondence should be addressed: Nicholas A. Graham, University of Southern California, Los Angeles, 3710 McClintock Ave., RTH 509, Los Angeles, CA 90089. Phone: 213-240-0449;

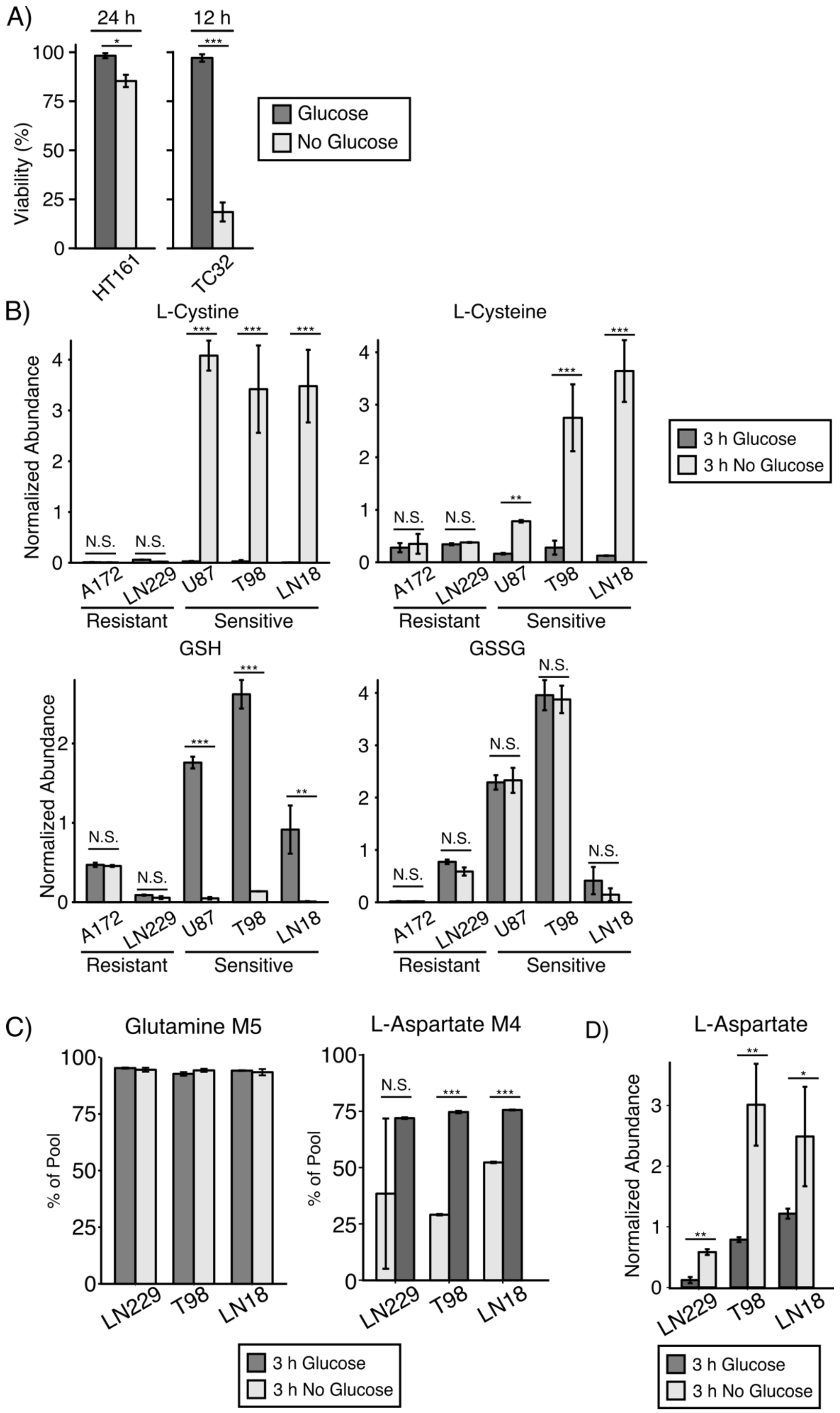
A)
24 h
12 h
Viability (%)
100
75
50
25
0
HT161
TC32
Glucose
No Glucose
B)
L-Cystine
L-Cysteine
GSH
GSSG
Normalized Abundance
N.S.
A172
LN229
U87
T98
LN18
Resistant
Sensitive
3 h Glucose
3 h No Glucose
C)
Glutamine M5
L-Aspartate M4
% of Pool
D)
L-Aspartate

**Supporting Figure 1. Accumulation of L-cystine and L-cysteine and depletion of reduced glutathione are metabolic markers of sensitivity to glucose deprivation.**

**A)** Two sarcoma cell lines were subjected to glucose deprivation, and viability was measured by trypan blue exclusion. HT161 exhibit strong resistance (<15% cell death at 24 h post glucose deprivation), whereas TC32 cells are highly sensitive to glucose deprivation (<25% viability at 12 h post glucose deprivation). * denotes $p < 0.05$, *** denotes $p < 0.001$ by Student's t-test.

**B)** Targeted metabolomics measurements in two glucose deprivation-resistant (A172, LN229) and three glucose deprivation-sensitive (T98, LN18, U87) cell lines confirmed that accumulation of L-cystine and L-cysteine, depletion of reduced glutathione (GSH), and unchanging levels of oxidized glutathione (GSSG) levels are hallmarks of sensitivity to glucose deprivation. N.S. denotes $p > 0.05$, ** denotes $p < 0.01$, *** denotes $p < 0.001$ by Student's t-test.

**C)** LC-MS metabolomics measurements of metabolites upon L-[U-$^{13}$C]-glutamine treatment. >90% of Glutamine was labeled after 3 h of treatment. Levels of L-Aspartate M4 increased upon glucose deprivation, indicating increased forward flux through the TCA cycle in all cell lines. N.S. denotes $p > 0.05$, *** denotes $p < 0.001$ by Student's t-test

**D)** LC-MS metabolomics measurements of metabolite abundance. Upon 3 h of glucose deprivation, L-Aspartate levels increased in all cell lines. * denotes $p < 0.05$, ** denotes $p < 0.01$ by Student's t-test.

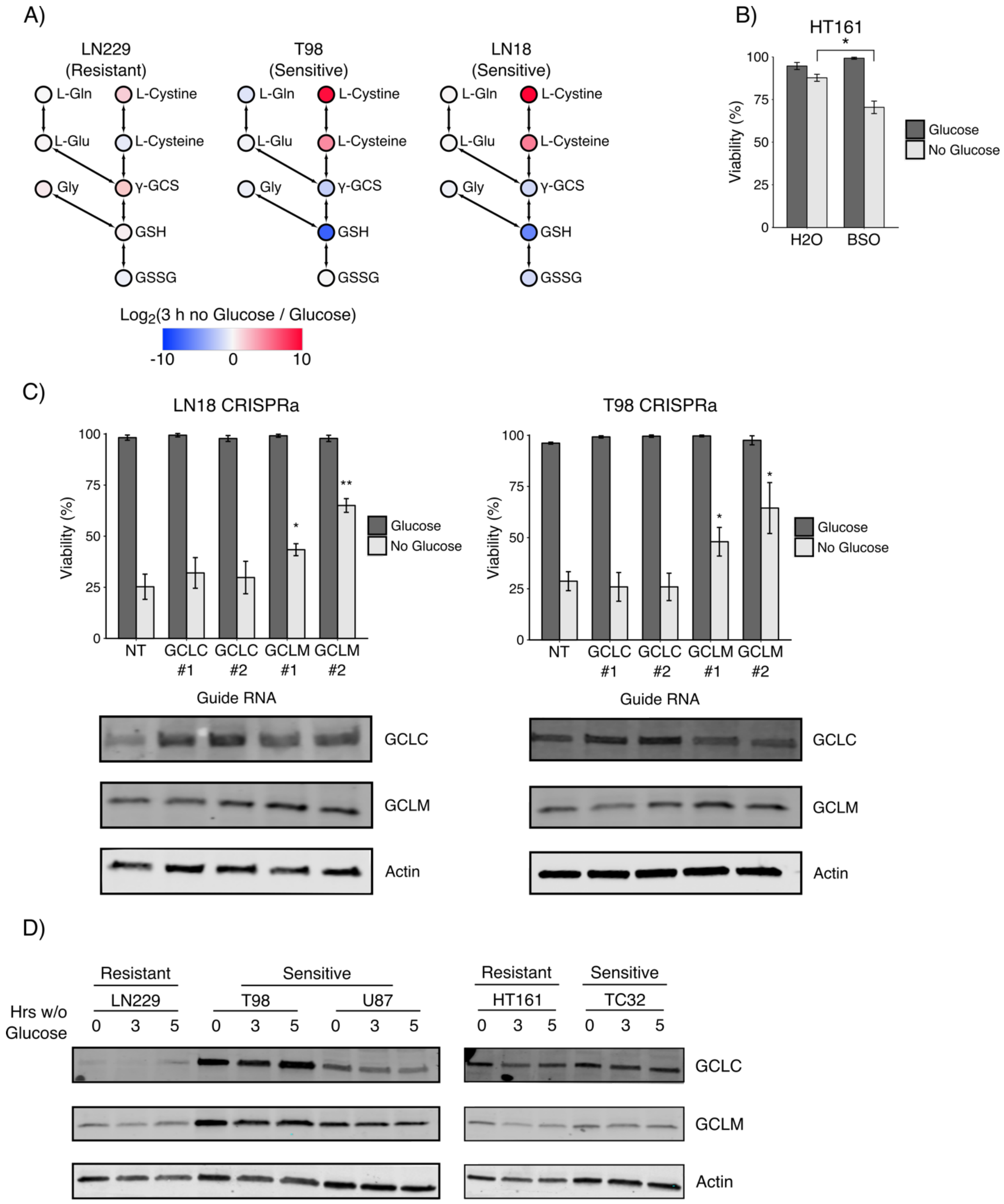


**Supporting Figure 2. GCL activity moderately regulates sensitivity to glucose deprivation.**

**A)** $Log_2$ fold change data from LN229, T98, and LN18 cells overlaid on glutathione metabolism pathway.

**B)** Treatment with the GCL inhibitor, buthionine sulfoximine (BSO, 500 μM), sensitized glucose deprivation-resistant HT161 sarcoma cells to glucose deprivation. Cells were treated with either solvent (water) or BSO in the presence of absence of glucose for 24 h, and viability was measured by trypan blue exclusion. * denotes $p < 0.05$ by Student's t-test (n=3).

**C)** Overexpression of GCLM but not GCLC increased resistance to glucose deprivation in sensitive LN18 and T98 cells. Cells were first infected with CRISPRa machinery (dCas9-VPR[1]) and then infected with a second vector carrying the indicated guide RNAs. Cells were then cultured by 8 h in the presence or absence of glucose, and viability was measured by trypan blue exclusion. * denotes $p < 0.05$, ** denotes $p < 0.01$ by Student's t-test (n=3). Western blotting with antibodies against GCLC, GCLM, and the equal loading control actin confirmed overexpression. NT, non-targeting control.

**D)** Expression of both GCLC and GCLM is stable upon glucose deprivation in both glucose deprivation-sensitive and -resistant cell lines. Glucose deprivation sensitive (T98, LN18, TC32) and resistant (LN229, HT161) cell lines were deprived of glucose for the indicated time points. Lysates were collected and western blots were analyzed with antibodies against GCLC, GCLM, and the equal loading control actin showed stable expression upon glucose deprivation in all cell lines.

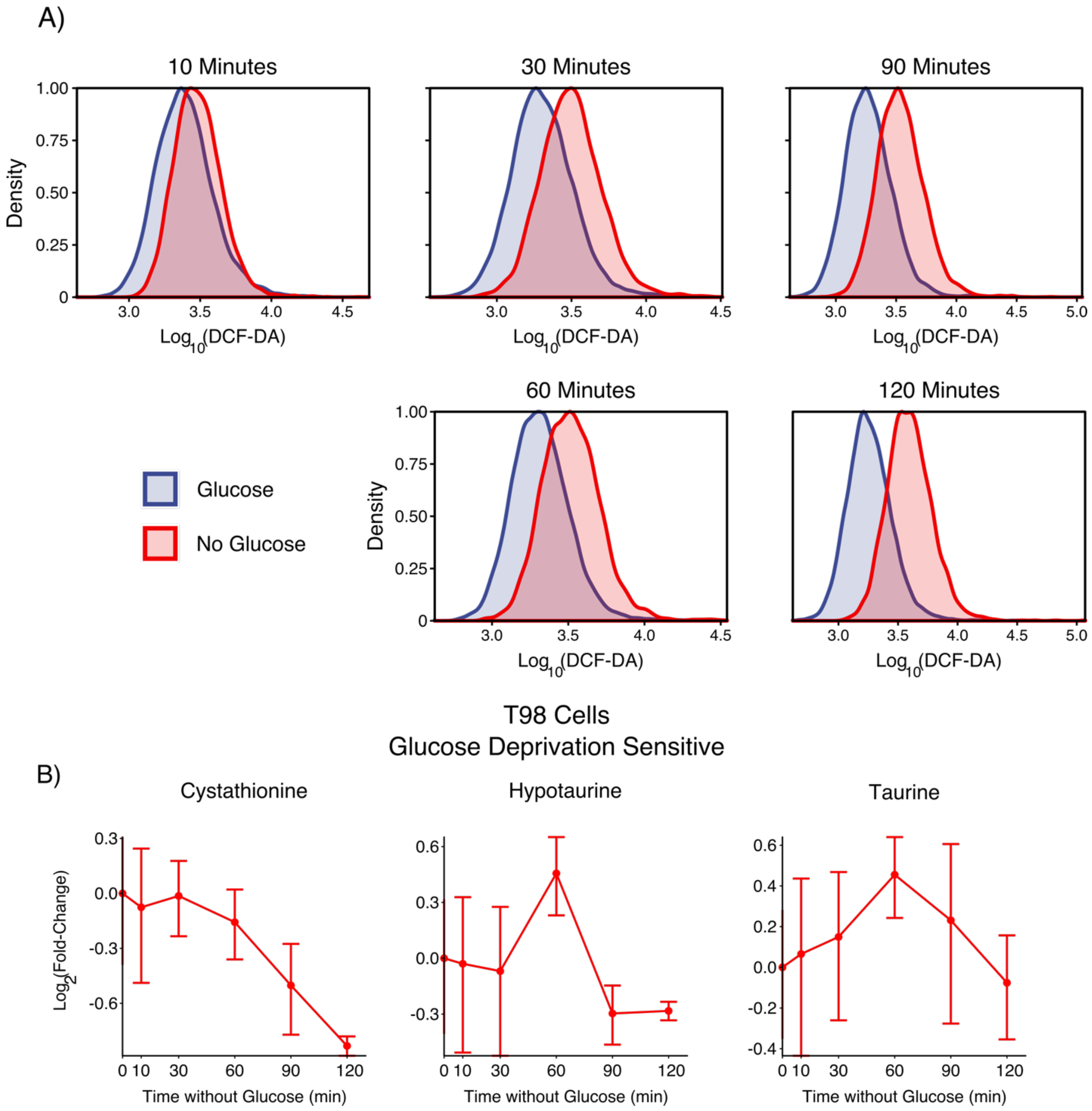


**Supporting Figure 3. Temporal measurements of reactive oxygen species and metabolite abundances upon glucose deprivation.**

**A)** Reactive oxygen species (ROS) accumulate upon glucose deprivation. The oxidation dependent fluorogen DCF-DA was used to measure ROS accumulation upon glucose deprivation. T98 cells were either starved of glucose (blue) or supplemented with fresh

DMEM with glucose (red) for the indicated times. Upon incubation with DCF-DA, cells were analyzed by flow cytometry.

**B)** Metabolite abundances of cystathionine, taurine, and hypotaurine do not increase upon glucose deprivation. Glucose deprivation-sensitive T98 cells were starved of glucose, and metabolite levels were measured by LC-MS metabolomics at the indicated time points. While levels of L-cysteine dramatically increased upon glucose deprivation (Fig. 3B), the alternative metabolic destinations of L-cysteine did not show substantial increases.

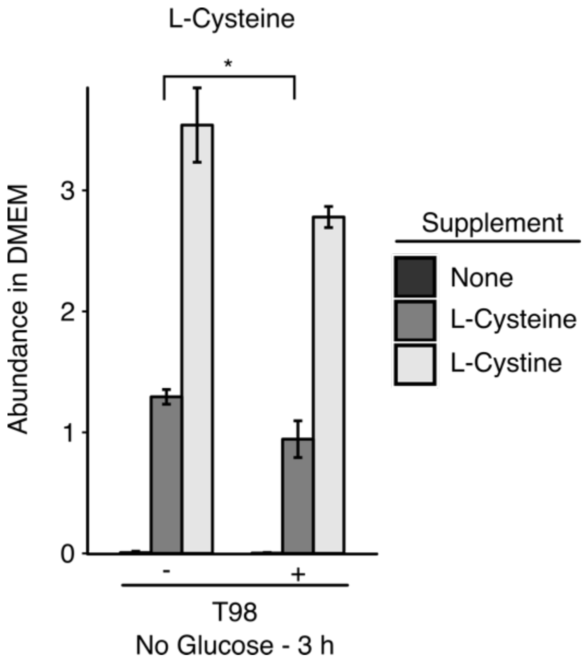


**Supporting Figure 4. L-cysteine is consumed by T98 cells.** L-cystine-free medium was supplemented with either nothing (control), L-cysteine (200 μM), or L-cystine (200 μM). T98 cells and control without cells were incubated for 3 h in each medium. Extracellular metabolites were extracted and measured using LC-MS metabolomics. L-cysteine was consumed by glucose deprived T98 cells. * denotes $p < 0.05$ by Student's t-test.

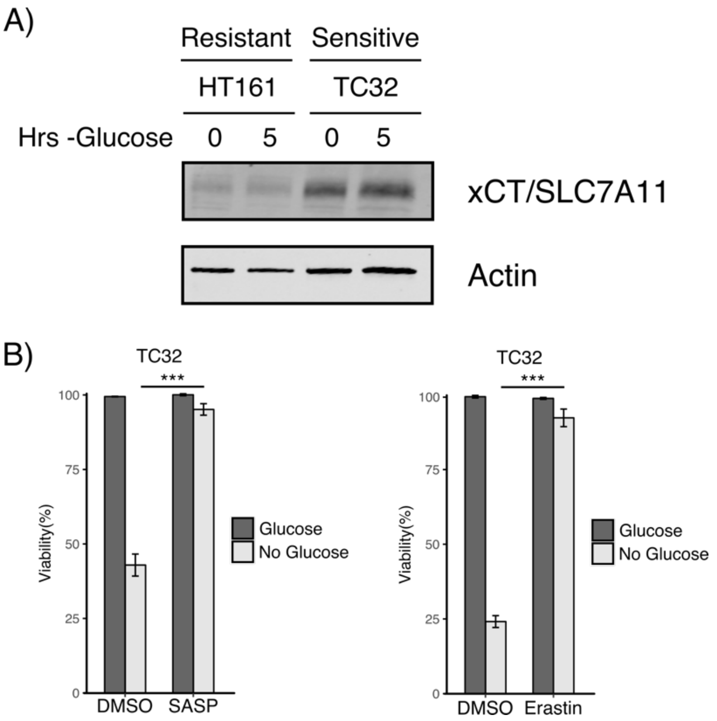


**Supporting Figure 5. L-cystine import from the glutamate/cystine antiporter, system $x_c^-$, promotes glucose deprivation-induced cell death.**

**A)** Expression of the specific light chain of system $x_c^-$, xCT/SLC7A11, correlated with sensitivity to glucose deprivation in sarcoma cell lines. The resistant cell line HT161 and the sensitive cell line TC32 were starved of glucose for the indicated time and then lysed. Expression of xCT/SLC7A11 was assessed by Western blotting. Actin was used as an equal loading control.

**B)** Pharmacological inhibition of xCT rescued sensitive cells from glucose deprivation-induced cell death. The sensitive sarcoma cell line TC32 was treated with either sulfasalazine (SASP, 500 μM) or erastin (10 μM) in the presence or absence of glucose for 8 h, and viability was assessed by trypan blue exclusion. *** denotes $p < 0.001$ by Student's t-test (n=3).

A) Glioblastoma

| Rank | Gene | Pearson Correlation | Permutation p-value | FDR q-value |
|---|---|---|---|---|
| 1 | *SLC2A1* | -0.439 | 0.015 | 0.220 |
| 2 | *HK3* | -0.410 | 0.029 | 0.220 |
| 3 | *LDHA* | -0.403 | 0.039 | 0.220 |
| 4 | *PGM1* | -0.397 | 0.060 | 0.220 |
| 5 | *FBP1* | -0.291 | 0.088 | 0.252 |
| 6 | *ALDH2* | -0.289 | 0.053 | 0.354 |
| 7 | *ALDH1B1* | -0.286 | 0.161 | 0.252 |
| 8 | *SLC16A4* | -0.277 | 0.103 | 0.252 |
| 9 | *GALM* | -0.272 | 0.096 | 0.252 |
| 10 | *SLC16A3* | -0.263 | 0.048 | 0.220 |

B) Ewing's Sarcoma

| Rank | Gene | Pearson Correlation | Permutation p-value | FDR q-value |
|---|---|---|---|---|
| 1 | *FBP1* | -0.662 | 0.005 | 0.084 |
| 2 | *G6PC* | -0.648 | 0.012 | 0.084 |
| 3 | *HK3* | -0.635 | 0.009 | 0.084 |
| 4 | *SLC16A3* | -0.563 | 0.018 | 0.084 |
| 5 | *BPGM* | -0.537 | 0.032 | 0.101 |
| 6 | *SLC2A1* | -0.530 | 0.040 | 0.110 |
| 7 | *SLC2A2* | -0.521 | 0.019 | 0.084 |
| 8 | *ENO2* | -0.489 | 0.049 | 0.120 |
| 9 | *ADPGK* | -0.488 | 0.111 | 0.244 |
| 10 | *PGAM2* | -0.486 | 0.028 | 0.101 |

C)

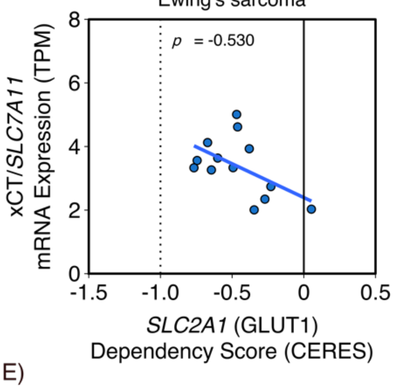


D)

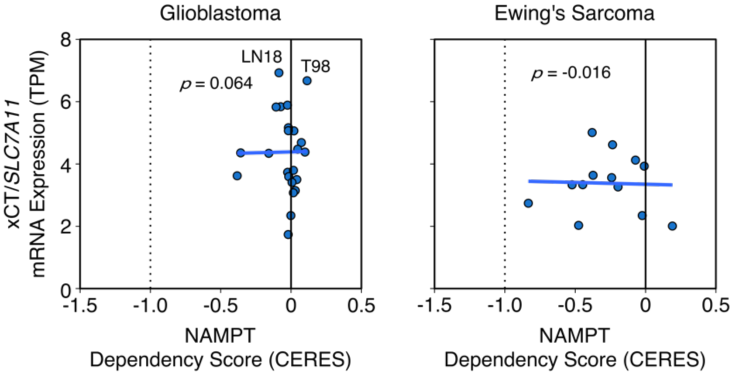


E)

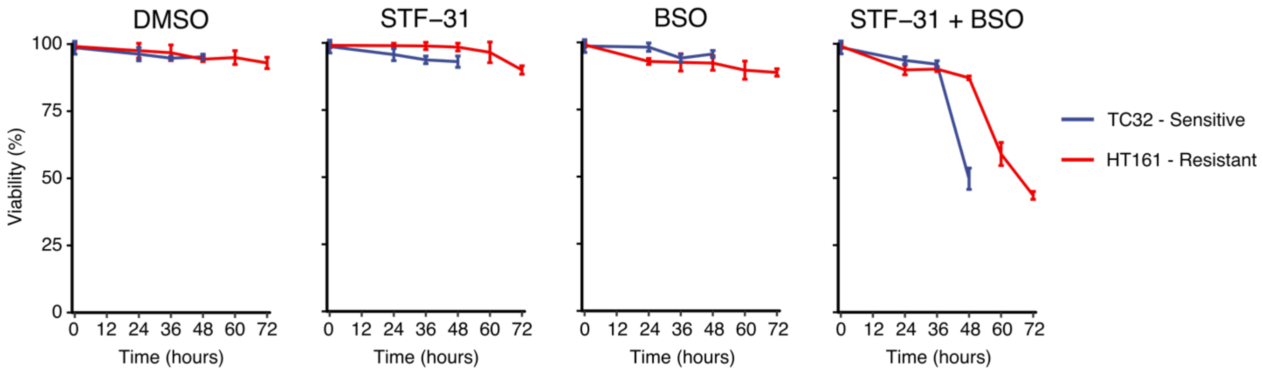


F)

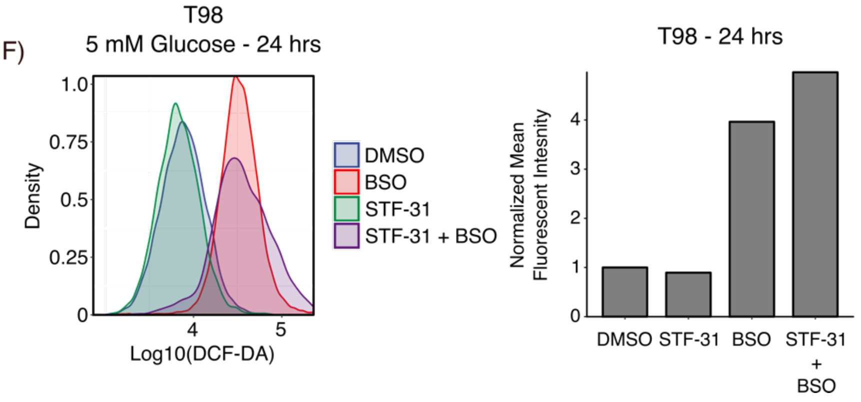

**Supporting Figure 6. Co-Inhibition of GLUT1 and GCL is synthetically lethal in glucose deprivation sensitive cell lines.**

**A,B)** Using data from the Cancer Dependency Map (DEPMAP[2]), glycolytic genes were filtered for expression in GBM or Ewing's Sarcoma and then ranked by their Pearson correlation coefficient between dependency (CERES score) and xCT expression. A negative CERES score indicates increased dependency. *SLC2A1* (GLUT1) was the top hit for GBM and the sixth highest hit for Ewing's sarcoma. Permutation p-values were assessed by permuting expression and dependency labels.

**C)** Ewing's sarcoma cells with increased xCT expression are more sensitive to *SLC2A1* (GLUT1) depletion. Data is from the Cancer Dependency Map (DEPMAP[2]) as in panel B.

**D)** NAMPT dependency does not correlate with xCT expression in glioblastoma and Ewing's sarcoma cell lines. Data is from the Cancer Dependency Map (DEPMAP[2]) as in panels A and B.

**E)** Glucose deprivation sensitivity correlated with response to combined STF-31 and BSO treatment. High sensitivity TC32 and low sensitivity HT161 sarcoma cells were cultured in 5 mM glucose and treated with STF-31 (12.5 µM), BSO (500 µM), or both drugs for the indicated amount of time. Glucose deprivation-sensitive TC32 cells died after 48 h of treatment, whereas resistant HT161 cells did not die until after 60 h of treatment.

**F)** Co-inhibition of GLUT1 and GCL induces ROS in T98 cells. Histograms showing the distribution of T98 cells treated with STF-31 (12.5 µM), BSO (500 µM), or both for 24 h. Combined treatment induced a 4-fold increase in the mean fluorescent intensity, whereas BSO alone induced a 3-fold increase in and STF-31 alone showed no increase compared to DMSO control.

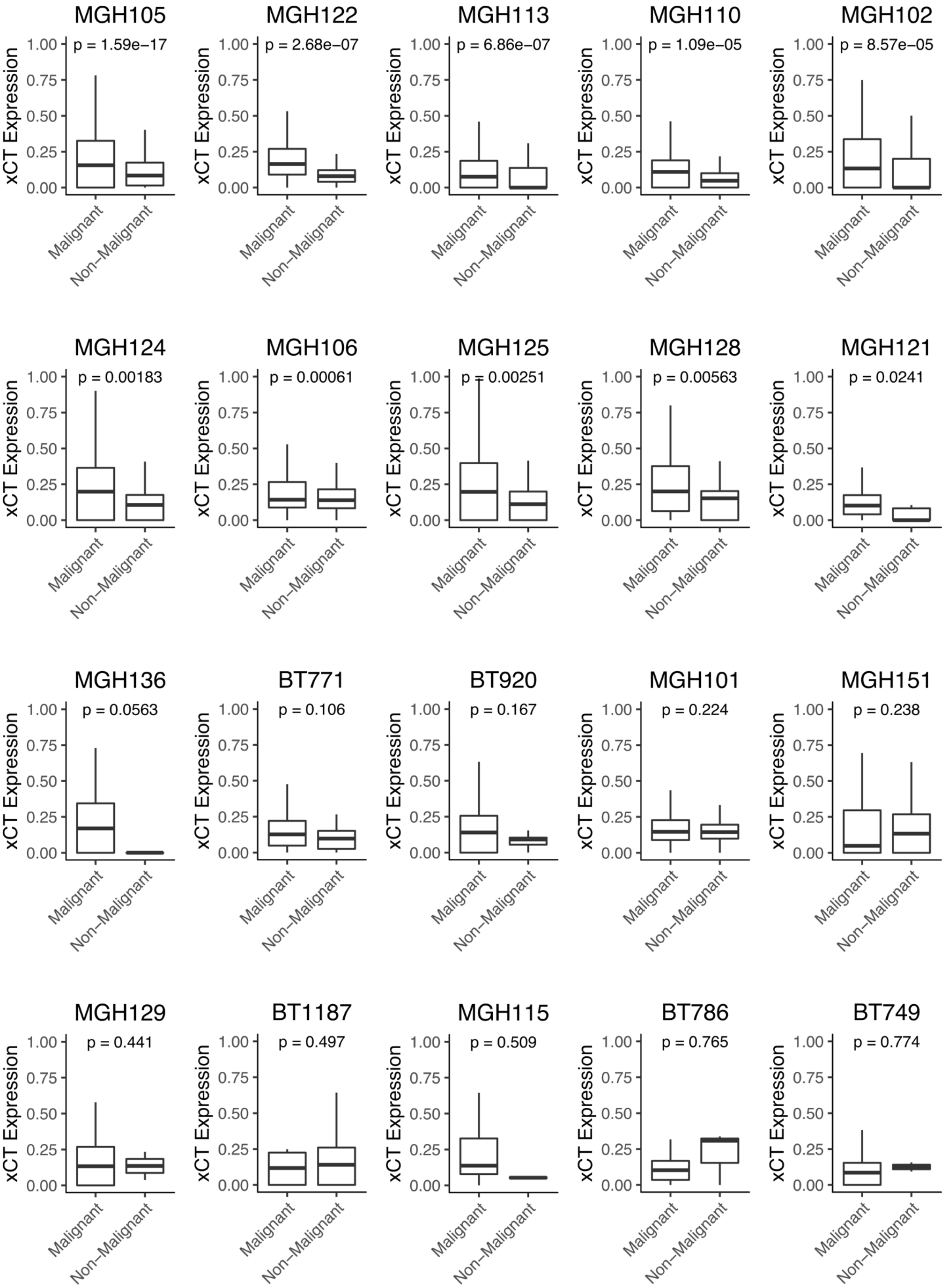
MGH105
p = 1.59e−17
MGH122
p = 2.68e−07
MGH113
p = 6.86e−07
MGH110
p = 1.09e−05
MGH102
p = 8.57e−05
MGH124
p = 0.00183
MGH106
p = 0.00061
MGH125
p = 0.00251
MGH128
p = 0.00563
MGH121
p = 0.0241
MGH136
p = 0.0563
BT771
p = 0.106
BT920
p = 0.167
MGH101
p = 0.224
MGH151
p = 0.238
MGH129
p = 0.441
BT1187
p = 0.497
MGH115
p = 0.509
BT786
p = 0.765
BT749
p = 0.774
xCT Expression
1.00
0.75
0.50
0.25
0.00
Malignant
Non−Malignant

**Supporting Figure 7. xCT expression is upregulated in a subset of glioblastoma patients.** Single-cell RNA-sequencing data was analyzed from adult and pediatric GBM patients[3] for xCT expression (logTPM). Data was filtered for tumors in which non-malignant cells were measured. 10 out of 20 patients exhibited significant increased expression of xCT in malignant cells compared to non-malignant cells as assessed using a Wilcoxon rank sum test.

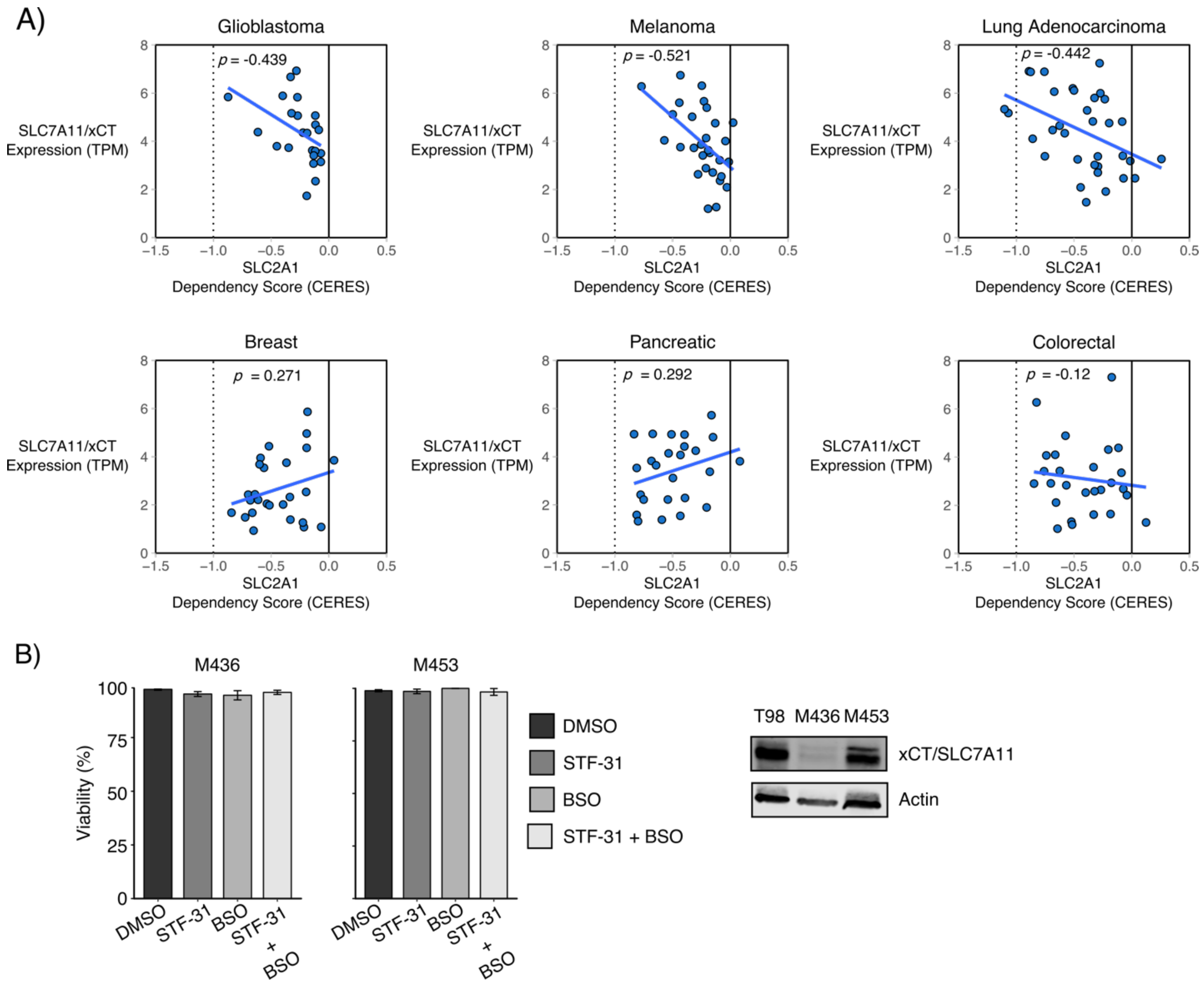


**Supporting Figure 8. Combined GLUT1 and GCL inhibition does not synergistically induce cell death in breast cancer cell lines.**

**A)** Cancer cell lines derived from certain tissues exhibit increased dependence on GLUT1 as xCT/SLC7A11 expression increases. Negative CERES scores (e.g. increased dependency) correlate with xCT expression in glioblastoma, melanoma, and lung adenocarcinoma cell lines, but not in breast cancer, pancreatic cancer, or colorectal cancer cell lines.

**B)** The breast cancer cell lines M453 (xCT high) and MB436 (xCT low) do not respond to STF-31, BSO, or combined STF-31 and BSO treatment after 72 h in 5 mM glucose.

Expression of xCT/SLC7A11 was assessed by Western blotting. Actin was used as an equal loading control.
